# Whole-brain serial-section electron microscopy in larval zebrafish

**DOI:** 10.1101/134882

**Authors:** David Grant Colburn Hildebrand, Marcelo Cicconet, Russel Miguel Torres, Woohyuk Choi, Tran Minh Quan, Jungmin Moon, Arthur Willis Wetzel, Andrew Scott Champion, Brett Jesse Graham, Owen Randlett, George Scott Plummer, Ruben Portugues, Isaac Henry Bianco, Stephan Saalfeld, Alex Baden, Kunal Lillaney, Randal Burns, Joshua Tzvi Vogelstein, Alexander Franz Schier, Wei-Chung Allen Lee, Won-Ki Jeong, Jeff William Lichtman, Florian Engert

## Abstract

Investigating the dense meshwork of wires and synapses that form neuronal circuits is possible with the high resolution of serial-section electron microscopy (ssEM)^1^. However, the imaging scale required to comprehensively reconstruct axons and dendrites is more than 10 orders of magnitude smaller than the spatial extents occupied by networks of interconnected neurons^2^—some of which span nearly the entire brain. The difficulties in generating and handling data for relatively large volumes at nanoscale resolution has thus restricted all studies in vertebrates to neuron fragments, thereby hindering investigations of complete circuits. These efforts were transformed by recent advances in computing, sample handling, and imaging techniques^1^, but examining entire brains at high resolution remains a challenge. Here we present ssEM data for a complete 5.5 days post-fertilisation larval zebrafish brain. Our approach utilizes multiple rounds of targeted imaging at different scales to reduce acquisition time and data management. The resulting dataset can be analysed to reconstruct neuronal processes, allowing us to, for example, survey all the myelinated axons (the projectome). Further, our reconstructions enabled us to investigate the precise projections of neurons and their contralateral counterparts. In particular, we observed that myelinated axons of reticulospinal and lateral line afferent neurons exhibit remarkable bilateral symmetry. Additionally, we found that fasciculated reticulospinal axons maintain the same neighbour relations throughout the extent of their projections. Furthermore, we use the dataset to set the stage for whole-brain comparisons of structure and function by co-registering functional reference atlases and *in vivo* two-photon fluorescence microscopy data from the same specimen. We provide the complete dataset and reconstructions as an open-access resource for neurobiologists and others interested in the ultrastructure of the larval zebrafish.

---

Pioneering studies in invertebrates established that synaptic-resolution wiring diagrams of complete neuronal circuits are valuable tools for relating a nervous system’s structure to its function^3-6^. Such resources can be combined with perturbations, activity maps, or behavioural assays to examine how signalling through neuronal networks transforms information from the environment into relevant motor outputs^5-10^. These studies benefited from the small size of the model organisms and stereotypy across individuals, which allow for complete ssEM of an entire individual or mosaicking of data from multiple individuals.

Vertebrate model nervous systems, on the other hand, are considerably larger and more variable. Consequently, ssEM of whole vertebrate neuronal circuits requires rapid computer-based technologies for acquiring, storing, and analysing many images from one animal. In many cases, anatomical data must be combined with other experiments on the same individual^11-13^ to define the relationship between structure, function, and behaviour. Because vertebrate nervous systems can vary substantially between individuals^14^, distilling a particular circuit’s structure may require its full description in multiple individuals. Applied to mammalian brains, this kind of analysis would require imaging very large volumes that are technically still out of reach (but see ref. 15). Thus, in mammals, ssEM has been confined to smaller and often partial reconstructions of circuits^16-23^. One strategy for capturing the brain-wide circuit underpinnings of behaviours is to generate whole-brain datasets with nanoscale resolution in smaller animals.

The larval zebrafish is an ideal system for this endeavour. It is a near-transparent vertebrate model organism that offers convenient optical access to its entire nervous system. Indeed, recent work has taken advantage of whole-brain calcium imaging in larval zebrafish^24-27^ and also ssEM examination of specific subregions^28-30^. Integrated with an established genetic toolkit and a variety of quantitative behavioural assays^28^, it is an excellent model organism for investigating the neuronal basis of behaviour^31^. In addition, its small size makes the larval zebrafish ideally suited for ssEM examination of even the complete brain.

Our goal was to develop a framework for ssEM of complete larval zebrafish brains at 5–7 days post-fertilisation (dpf), which coincides with the emergence of complex behaviours such as prey capture^32^ and predator avoidance^33^. Attempts to apply existing methods to the larval zebrafish were impeded by two obstacles: First, skin and membranes covering the brain^34^ prevent high-quality tissue preservation by whole-fish immersion alone. Second, consistent ultrathin sectioning is difficult to achieve in samples containing heterogeneous tissues but imperative for reconstructing three-dimensional structure from a series of two-dimensional sections.

To preserve ultrastructure across the complete brain with minimal damage, we developed a fine dissection technique to remove skin and membranes from the dorsum (see Methods; Extended Data Fig. 1a–c). Following dissection, traditional EM protocols (Extended Data Fig. 1d–h) resulted in high-quality tissue preservation throughout the brain (Extended Data Fig. 1i).

Consistent sectioning perpendicular to the majority of axon and dendrite paths is preferable for ease and reliability in reconstructing neuronal morphology. Therefore, we sectioned perpendicular to the long (anterior-posterior) axis of one 5.5dpf larval zebrafish, which required ∼2.5× more sections than the horizontal orientation (dorsal-ventral). This was made possible by customising an automated tape-collecting ultramicrotome^35^ to expand its tape-carrying capacity (Extended Data Fig. 2a–c) such that all sections could be collected on one continuous tape stretch. Furthermore, sectioning in test samples revealed that errors and loss occurred primarily when the composition of the sample changed dramatically (e.g., the border of embedded tissue and empty resin). We overcame this sectioning difficulty by embedding the sample in a low-viscosity epoxy resin within a surrounding support tissue of mouse cerebral cortex (Extended Data Fig. 1g–h, Extended Data Fig. 2f). This approach generated a stable library of sections on silicon wafers (Extended Data Fig. 2d–e) that could be imaged multiple times and at different resolutions.

Overview images were acquired to survey all sections (Extended Data Fig. 3; Supplementary Video 1), resulting in a stack of 17963 images spanning 1.02×10^10^μm^3^, consisting of 3.01×10^11^ voxels, and occupying 310 gigabytes of computer storage. In total, 17963 × ∼60nm-thick sections were collected from 18207 attempted, with 244 lost (1.34%; Extended Data Fig. 3d top) and 283 containing partial regions of the tissue (1.55%; Extended Data Fig. 3d middle; Extended Data Fig. 5). There were no adjacent losses and 5 instances of adjacent lost-partial or partial-partial events (0.03%; Extended Data Fig. 3d bottom). From visualizations of this low-resolution data, we confirmed that our approach enabled stable serial sectioning through a millimetre-long region spanning from myotome 7 to the anterior-most larval zebrafish structures—encompassing the anterior spinal cord and the entire brain (Extended Data Fig. 3b–c; Supplementary Video 2).

We next selected sub-regions within this imaging volume to capture areas of interest at higher resolutions using multi-scale imaging^35,36^ (Fig. 1). We first performed nearly isotropic EM imaging by setting lateral resolution to match section thickness over the anterior-most 16000 sections (Fig. 1a–f; Supplementary Video 3). All cells are labelled in ssEM, so this volume offers a dense picture of the fine anatomy across the anterior quarter of the larval zebrafish including brain, sensory organs (e.g., eyes, ears, and olfactory pits), and other tissues. Furthermore, this resolution of 56.4×56.4×60nm^3^/vx is ∼500× greater than that afforded by diffraction-limited light microscopy. The imaged volume spanned 2.28×10^8^μm^3^, consisted of 1.12×10^12^ voxels, and occupied 2.4 terabytes (TB). In this data, one can reliably identify cell nuclei and track large-calibre myelinated axons (Fig. 1b–c,e–f; Supplementary Video 4). To further resolve densely packed neuronal structures, a third round of imaging at 18.8×18.8×60nm^3^/vx was performed to generate a high-resolution atlas specifically of the brain. The resulting image volume (Fig. 1d,g– h) spanned 12546 sections, contained a volume of 5.49×10^7^μm^3^, consisted of 2.36×10^12^ voxels, and occupied 4.9TB. Additional acquisition at higher magnifications was used to further inspect regions of interest, to resolve finer axons and dendrites, and to identify synaptic connections between neurons (Fig 1i–k). All images across sections and scales were co-registered form a coherent multi-resolution ssEM dataset (Supplementary Video 3).

**Figure 1:**
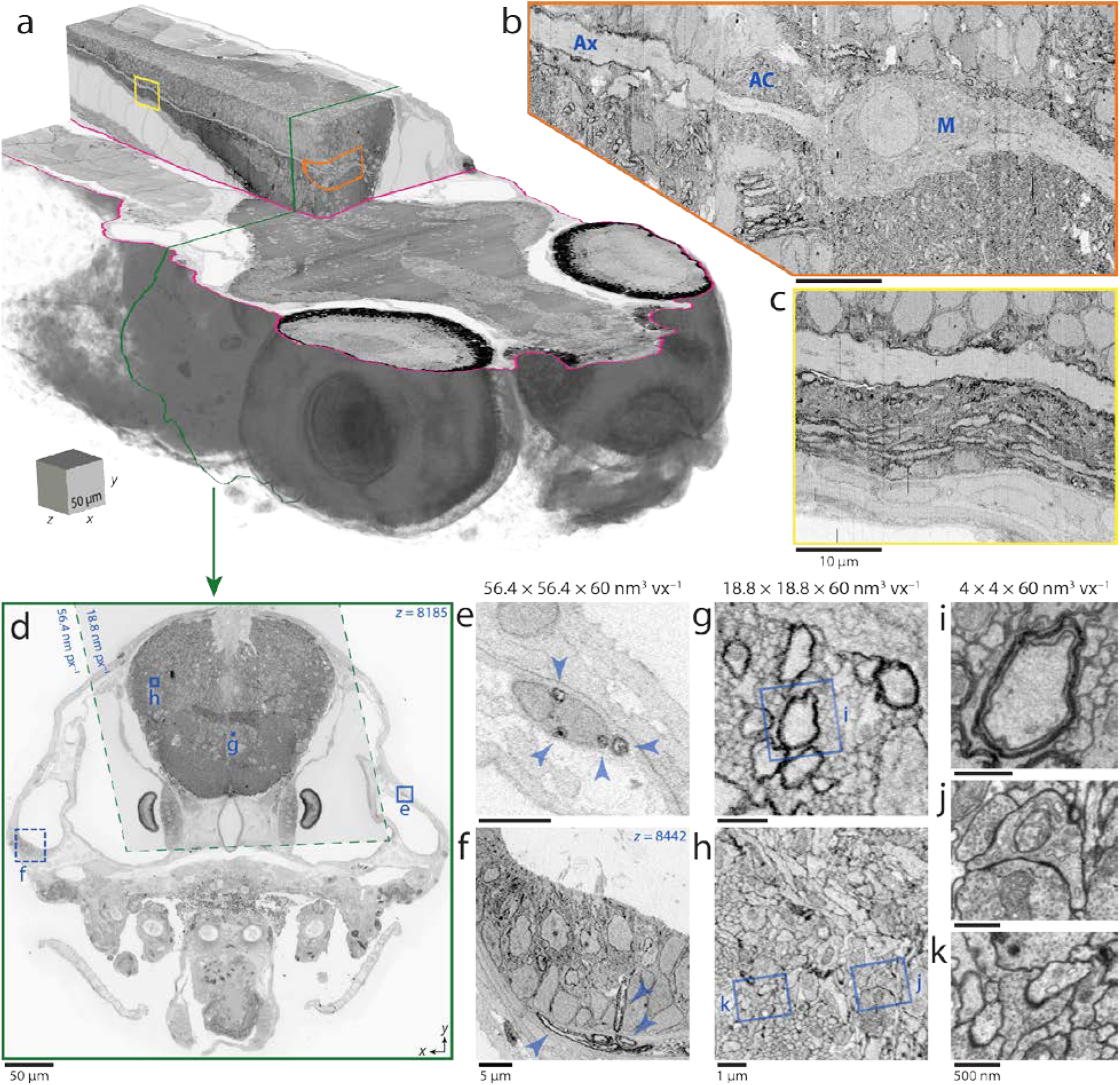
Targeted, multi-scale ssEM of the larval zebrafish brain. **a**, Volume rendering of and reslices through 16000 aligned serial micrographs that form a nearly isotropic (56.4×56.4×60nm^3^vx^−1^) image volume containing the anterior quarter of a larval zebrafish. **b–c**, Flattened contour reslice images reveal brain features visible with nearly isotropic resolution. **b**, The Mauthner cell (M), axon cap (AC), and axon (Ax) surrounded by other neurons, myelinated structures, and neuropil. **c**, Posterior extension of the Mauthner axon. **d**, Micrographs from a single section acquired in separate rounds of imaging. All larval zebrafish tissue in 16000 sections was acquired at nearly isotropic resolution (solid border, **e–f**). The brain alone in 12546 sections was acquired at higher (18.8×18.8×60nm^3^vx^−1^) resolution (dashed border, **g–h**). **e–f**, Large-caliber myelinated structures (arrowheads) are identifiable in nearly isotropic micrographs that include peripheral nerves (**e**) and sensory organs such as the ear (**f**, image corresponds to the dashed region in **d** from a nearby section). **g–h**, Many neuronal processes including myelinated fibers can be segmented from higher resolution micrographs. **i–k**, Additional rounds of targeted re-imaging can be performed to distinguish finer neuronal structures and the connections between them. Scale box: **a**, 50×50×50μm^3^. Scale bars: **b**–**c**, 10μm; **d**, 50μm; **e**–**f**, 5μm; **g**–**h**, 1μm; **i**–**k**, 500nm.

With a framework in place for producing whole-brain ssEM data (Extended Data Fig. 6), we next tested the feasibility of identifying individual neurons and fluorescently defined regions across imaging modalities, as done for smaller volumes previously^11-13^. To accomplish this, we matched the positions of nuclei in the ssEM dataset with those in two-photon fluorescence of a genetically encoded calcium indicator acquired from the same animal. The common structural features between the two image sets enabled identification of the same neurons in both datasets (Extended Data Fig.7; Supplementary Video 5). Furthermore, this revealed the imaging conditions, labelling density, and structural tissue features necessary for reliable matching across imaging modalities. It was difficult in regions where fluorescence signal was low (Extended Data Fig. 7e), where many cells were packed closely together (Extended Data Fig. 7f), and where progenitors add new neurons in between light microscopy and preparation for ssEM (Extended Data Fig. 7g). Improving the light-level data with specific labelling of all nuclei and faster light-sheet or other imaging approaches should greatly improve the ease and accuracy of matching neuron identity across modalities. This ability to assign neuron identity across imaging modalities demonstrates proof-of-principle for the potential integration of rich neuronal activity maps with subsequent whole-brain structural examination of functionally characterized neurons and their networks. By following a similar process, we were also able to transform the Z-Brain reference atlas^37^ and Zebrafish Brain Browser^38^, expandable open-source zebrafish references containing a variety of molecular labels, to seamlessly integrate into the multi-resolution ssEM dataset (Extended Data Fig. 6; Extended Data Fig. 8).

We next tested the general applicability of this dataset for reconstructions of selected neurons. First, we reconstructed a peripheral lateral line afferent neuron that innervates hair cells in a dorsal neuromast sensory organ and projects to the hindbrain with its soma residing in the posterior lateral line ganglion (Fig. 2a–e; Supplementary Video 6). By re-imaging this neuromast at higher resolution (4×4×60nm^3^/vx), we identified synapses connecting hair cells and this primary sensory neuron (Fig. 2b–c) while improving reconstructions in unmyelinated regions in the periphery (Fig. 2d). We also reconstructed myelinated motor neurons, including several in the spinal cord (Fig. 2f), that directly contacted muscle. Further, myelinated neurons could also be identified and annotated within the brain. These reconstructions highlight the utility of multi-resolution ssEM for reassembling morphologies of various large-calibre myelinated neurons from sensory inputs, throughout the brain, and back to peripheral innervation of muscle.

**Figure 2:**
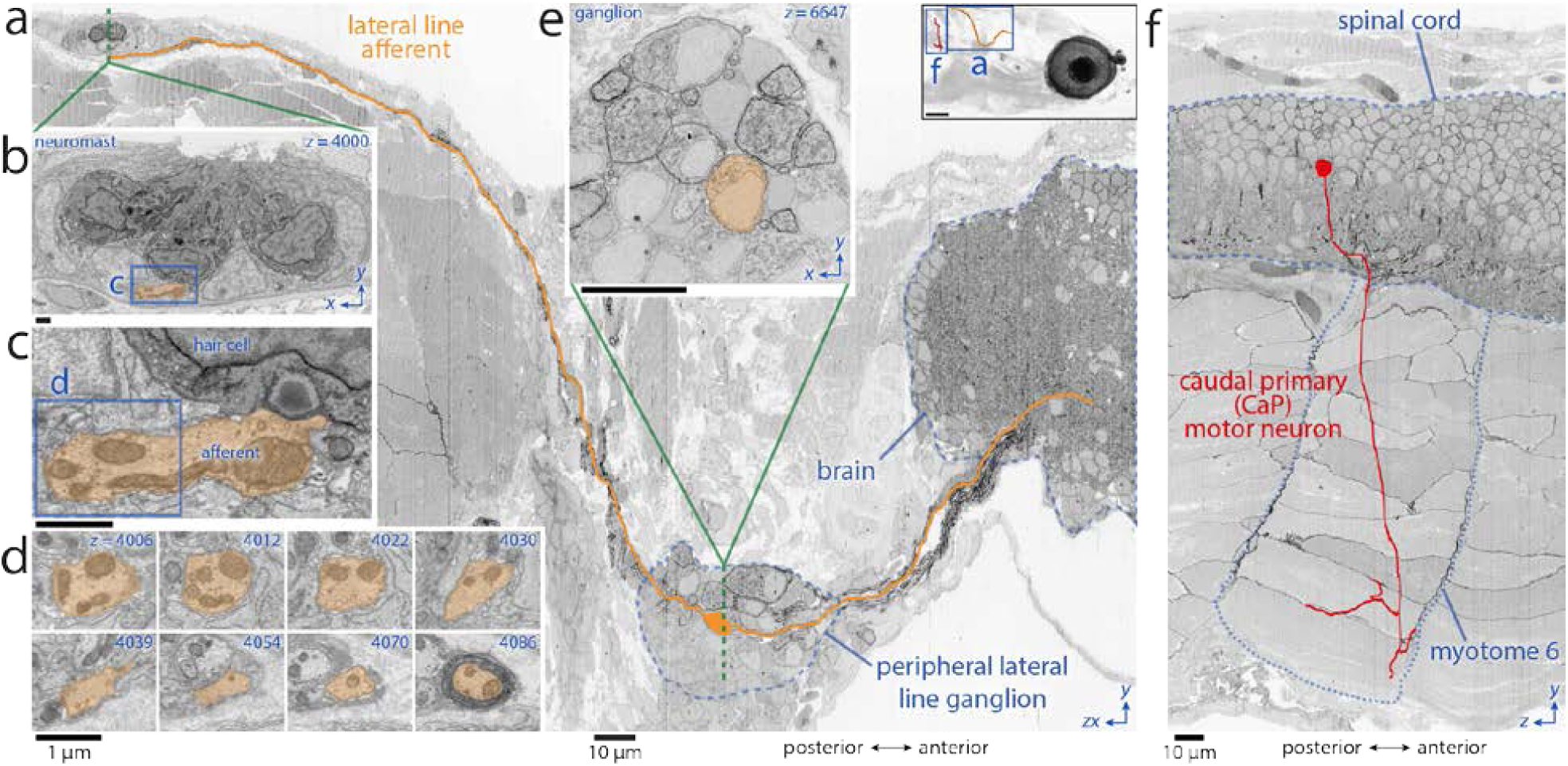
Neuronal morphology reconstructions capturing sensory input and motor output. **a**, The path of a bipolar lateral line afferent neuron (orange) is tracked from a synaptic contact with a neuromast hair cell (**b**–**d**) through the peripheral lateral line ganglion (**e**) into the hindbrain, as revealed by a contour reslice through ∼5000 serial sections. Inset at top right is a lateral view of reconstructions in this figure, illustrating their position. **b**, Cross-section through the dorsal neuromast innervated by the highlighted afferent. **c**, Magnified ribbon synapse connecting a neuromast hair cell to the highlighted afferent. **d**, Series of micrographs showing the afferent’s exit from the neuromast and myelination. **e**, Cross-section through the peripheral lateral line ganglion showing the myelinated soma of the afferent. **f**, The position of a caudal primary (CaP) motor neuron in the spinal cord and its innervation of myotome 6 projected onto a reslice through ∼2200 serial sections. Scale bars: **a** top right inset, 100μm; **a**, **e**–**f**, 10μm; **b**–**d**, 1μm.

This ability to reconstruct individual neurons throughout the entire brain allows for the generation of complete circuit descriptions, a necessity for identifying the pathways that underlie entire behavioural programs. To exploit this potential opportunity, we produced a ‘projectome’ consisting of all the myelinated axons within this larval zebrafish’s brain (Fig. 3a–b; Supplementary Video 7; see Methods). We reconstructed 2589 myelinated axon segments along with many of their somata and dendrites to yield 39.9cm of combined axon and dendrite length. We could easily follow 834 myelinated axons comprising 30.6cm back to their somata. For the remaining axon segments (9.3cm), it was more difficult to identify their somata because of long unmyelinated stretches. The longest individual axon reconstructed, a trigeminal ganglion sensory afferent, was followed for 1.2mm and extended from its sensory terminals in the anterior skin surface to its termination field in the brain.

**Figure 3:**
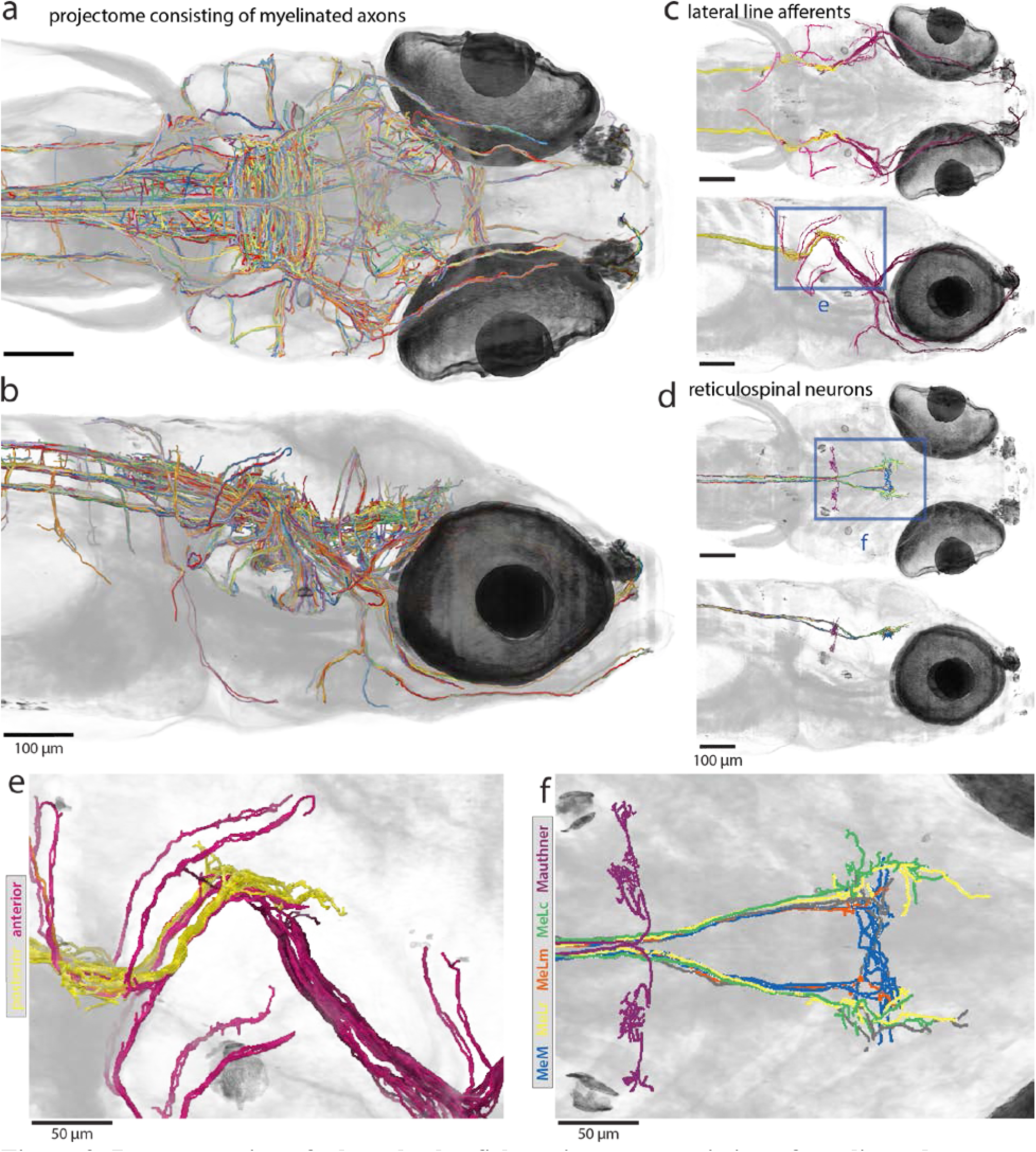
Reconstruction of a larval zebrafish projectome consisting of myelinated axons. **a–b**, Dorsal (**a**) and lateral (**b**) views of myelinated axon reconstructions manually extracted from the multi-resolution ssEM dataset. Colors are randomly assigned. **c**, Dorsal (top) and lateral (bottom) views of reconstructed lateral line primary sensory neurons. Anterior (purple) afferents were identified by their innervation of neuromasts in the periphery, where darker purple indicates more anterior. Posterior (yellow) afferents were identified by their presence in the posterior lateral line nerve. **d**, Dorsal (top) and lateral (bottom) views of a reticulospinal neuron subset including the Mauthner cell and nucleus of the medial longitudinal fasiculus (nucMLF) neurons. Cell identity was assigned by position and morphology for the Mauthner cell (purple) and the four canonical nucMLF neurons: MeLc (green), MeLr (yellow), MeLm (orange), and MeM (blue). **e**, Magnified view from **c** reveals topographically arranged afferent hindbrain axons such that those innervating more anterior neuromasts sit ventrolateral to those innervating more posteriorly located neuromasts. **f**, Magnified view of Mauthner cell and nucMLF neuron morphologies from **d**. Note the apparent bilateral symmetry in **c**–**f**. Scale bars: **a**–**d**, 100μm; **e**–**f**, 50μm.

The resulting projectome included 94 lateral line afferents that innervated 41 neuromasts through their ganglia to the hindbrain (Fig. 3c,e). Hindbrain arborisations of lateral line afferents were, as expected, topographically arranged such that those innervating more anterior neuromasts were ventrolateral to those innervating more posteriorly located neuromasts^39^ (Fig. 3e). These reconstructions reveal a striking bilateral symmetry in the lateral line system (Supplementary Video 8). Only one neuromast and its afferents did not have counterparts on the other side (Supplementary Video 8, orange sphere).

The projectome also included a significant fraction of the reticulospinal neuron population, which sends myelinated axons to the spinal cord from the midbrain and hindbrain. Similar to what we observed in the lateral line system, these neurons exhibit bilateral symmetry (Fig. 4a, red label). Additionally, we were able to identify individual reticulospinal neurons by their known positions and morphologies^40,41^ (Fig. 3d,f; Fig. 4a; Extended Data Fig. 9i; Supplementary Video 9). This afforded us an opportunity to precisely examine the extent of symmetry in this system.

**Figure 4:**
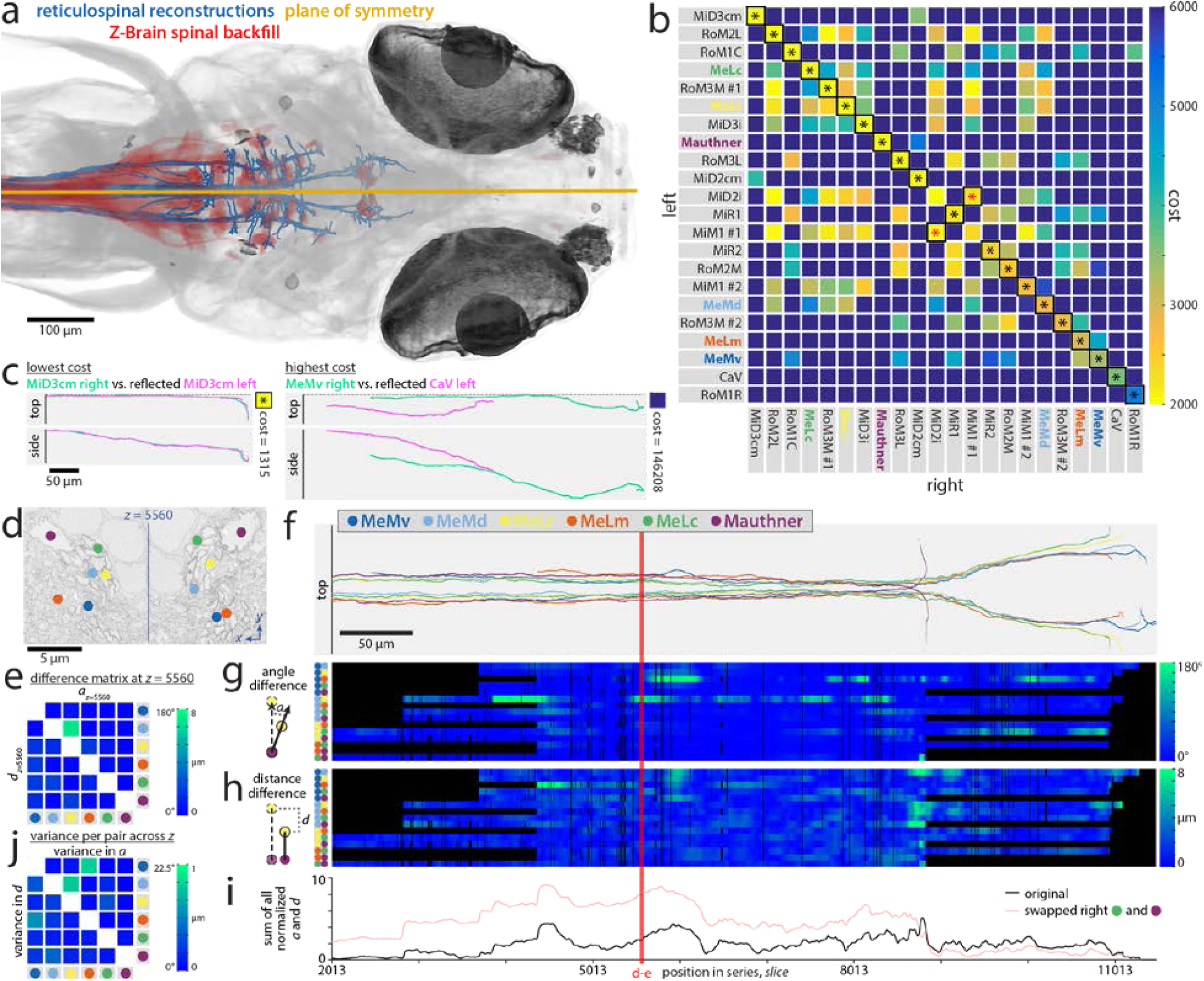
Symmetry in myelinated axons of the reticulospinal system. **a**–**c**, Three-dimensional (3D) symmetry analysis for 44 identified neurons (22 per side) whose axon traveled in the ∼30μm-diameter medial longitudinal fasciculus (MLF). **a**, Dorsal view of identified reticulospinal axons overlaid by the co-registered Z-Brain spinal backfills label. The plane of symmetry was fit from all identified axons that formed part of the MLF and used in subsequent analyses. **b**, Pairwise bilateral symmetry comparisons in 3D. Each axon on one side was compared to the reflection of every axon on the other side across the symmetry plane. Costs were calculated as the sum of the Euclidian distances between points matched by a dynamic time warping algorithm, then normalized by the number of matches and penalized by any unmatched regions (see Extended Data Fig. 9 and Methods). The matrix is ranked by cost between left-right homologs along the diagonal (top-left to bottom-right). Asterisks indicate the results of a globally optimal pairwise assignment. Though the analysis lacked any bias based on neuron identity, in all but one pair (red asterisks), the axon on the left was assigned to its homolog on the right. Low costs observed in off-diagonal elements highlight interesting similarities between the different neuron types. **c**, Top and side views of pairs resulting in the lowest (left) and highest (right) costs. **d**–**j**, Two-dimensional (2D) symmetry analysis of neighbour relations between axon pairs in one bundle with respect to the reflected pair from the other side for 6 left-right pairs of identified neurons (12 in total). Axon identity color code in **f** applies to all **d**–**j** panels. **d**, In cross-section, most axons within the left and right MLF appear to share mirror symmetric relative positions. Line indicates midline. **e**, Angle and distance differences for all 15 pairs in a single cross-section (slice 5560, indicated by red line in **f**–**i**), where the vector connecting a pair on one side is compared to the reflected vector between the same pair from the opposite side (see Methods). **f**, Top view of 6 identified axon pairs for which neighbor relations were examined. **g**– **h**, Linearizing the angle (**g**) and distance (**h**) differences between all pairs reveals that positional arrangements are mirror symmetric with their contralateral counterparts over long stretches of the MLF. Additionally, the neighbour relations return to a state of symmetry even after local perturbations. Black regions indicate where reconstruction data does not exist for a given pair. **i**, Summing normalized angle and distance differences shows particular regions (peaks) with neighbour relation perturbations, such as the entry of a new axon into the bundle. Swapping the identity of one pair of axons (MeLc and Mauthner) on one side results in a consistent near-doubling of this sum over for all cross-sections in which they both exist. **j**, Calculating the angle and distance difference variance across all cross-sections identifies a few pairs with weak neighbor relations. Scale bars: **a**, 100μm; **c**,**f**, 50μm; **d**, 5μm.

From the projectome, we selected 44 identified reticulospinal neurons (22 on each side) whose myelinated axons form the medial longitudinal fasciculus (MLF)^41^ to quantitatively analyse the degree of bilateral symmetry in specific projections. We started by investigating whether the myelinated axons in the MLF of one hemisphere are symmetric in their shape and position to that of the homologous neurons on the contralateral side. Notably, computing a globally optimal pairwise assignment (Fig. 4b, asterisks) resulted in excellent matching of left-right homologs (Fig. 4b, diagonal) in all but one pair (Fig. 4b, red asterisks; Extended Data Fig. 9d).

In addition, we also inquired about the possibility of bilateral symmetry in the neighbour relations between myelinated axons within the nerve bundle. In cross-sections, we observed that axons appeared to be situated in relatively similar positions within the left and right bundle, suggesting that bilateral symmetry could also extend to the neighbour relations within the MLF (Fig. 4d, compare relative positions across line). We developed quantitative metrics for comparing the spatial relations between an identified axon and its neighbours within the bundle on one side of the brain versus the other side (see Methods; Fig. 4e,g-h; Supplementary Video 10). We selected 6 left-right pairs of identified neurons (12 axons in total; Fig. 4f) and calculated the positional relationships of all the 15 (i.e., 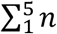) pairwise combinations between neighbouring axons in one bundle with respect to the reflected pair from the other side (Extended Data Fig. 9e-j). We found that the positional arrangements of these axons are mirror symmetric with their counterparts on the contralateral side. This symmetry in neighbour relations was evident over long stretches of the nerve bundles (Fig. 4f-j; Supplementary Video 10). Moreover, the axon neighbour relations returned to a state of symmetry even in the face of local perturbations such as the entry of a new axon into the bundle (Fig. 4f-i; Supplementary Video 10). Similar relationships were seen in the larger set of 44 reticulospinal axons (Extended Data Fig. 9i-k).

Although these axons originate from stereotyped positions in the midbrain and hindbrain, we expected that their positions would become progressively more scrambled in the same way neighbour relations of axons in peripheral mammalian nerves change with distance and show no mirror symmetry^14^. Were that the case, the configuration of the axons in the two bundles should become considerably less symmetric as they travel further posterior toward the spinal cord. The fact that this kind of randomization in position does not occur raises the possibility that axon bundles preserve some positional information along their course. Such an idea has a precedent in the ribbon-shaped optic nerve in certain kinds of fish, but was thought to be a special feature of this structure^42^.

This work suggests that stereotyped neighbour relations may be a general feature of fasciculated central nerve tracts. Interestingly, not all axons seemed to obey this symmetry as consistently as others (Fig. 4j). It is intriguing that in mammals—where axon branches reorganize in development via activity dependent mechanisms—such symmetry is not known to occur. Perhaps fish have both plastic and hardwired axons in their central nervous system whose development is guided by distinct mechanisms.

Here we demonstrate the feasibility of reconstructing neurons in whole-brain ssEM datasets acquired from larval zebrafish. We illustrate the utility of multi-scale imaging for reducing both imaging time and data storage requirements. We show that imaging of section libraries on wafers allows repeated re-imaging at higher resolutions, an option not available with block-face ssEM methods. Furthermore, continued development of faster EM technologies^43^ will hasten the re-imaging process and permit whole-brain studies in a fraction of the time required here.

Finally, the datasets presented here are not limited to analyses of the nervous system and can easily be extended to examine the structure of other organ systems (Extended Data Fig. 10), such as musculoskeletal, cardiac, intestinal, and pancreatic tissues. The data we generated can, thus, be used as an anterior larval zebrafish reference atlas and is available as an online resource (http://hildebrand16.neurodata.io/) for study by the scientific community.

## Methods

### Animal care

Adult zebrafish (*Danio rerio*) for breeding were maintained at 28°C on a 14hr:10hr light:dark cycle following standard methods^44^. The *Tg(elavl3:GCaMP5G)a4598* fish line^24^ used in this study was of genotype *elavl3*:GCaMP5G^+/+^; nacre (*mitfa*^−/−^), conveying nearly pan-neuronal expression of the calcium indicator GCaMP5G^45^ and increased transparency due to the nacre mutation^46^. The larval zebrafish described in this study were raised in filtered fish facility water^44^ until 5–7 days post-fertilization (dpf).

Mice from which support tissue was collected had been previously euthanized for unrelated experiments. Only unused, to-be-discarded tissue from these studies was harvested to serve as support tissue.

The Standing Committee on the Use of Animals in Research and Training of Harvard University approved all animal experiments.

### Two-photon laser-scanning microscopy

Larval zebrafish were immobilized by immersion in 1mg mL^−1^ α-bungarotoxin (Invitrogen) and mounted dorsum-up in 2% low-melting-temperature agarose in a small dish containing a silicone base (Sylgard® 184, Dow Corning). Upon agarose hardening, E3 solution (5mM NaCl, 0.17mM KCl, 0.33mM CaCl_2_, and 0.33mM MgSO_4_) was added to the dish. *In vivo* structural imaging of *elavl3*-driven nearly pan-neuronal GCaMP5G signal was conducted with a custom-built two-photon microscope with a Ti:Sapphire laser (Mai Tai®, Spectra-Physics) excitation source tuned to 800nm. Frames with a 764.4×509.6μm^2^ field-of-view size (1200×800px^2^) were acquired at 1μm intervals (0.637×0.637×1μm^3^vx^−1^) at ∼1 Hz with a scan pattern of four evenly spaced, interlaced passes, similar to a previously reported scanning approach^26^. A low-noise anatomical snapshot of complete brain fluorescence was captured in 300 planes, each the sum of 50 single frames. All light-based imaging was performed without any intentional stimulus presentation.

### Dissection and EM tissue preparation

Any aqueous solutions were prepared with water passed through a purification system (typically Arium 611VF, Sartorius Stedim Biotech).

Larval zebrafish embedded in low-melting-temperature agarose were introduced to a dissection solution (64mM NaCl, 2.9mM KCl, 10mM HEPES, 10mM glucose, 164mM sucrose, 1.2mM MgCl_2_, 2.1mM CaCl_2_, and pH 7.5; ref. ^47^) containing 0.02% (w/v) tricaine mesylate (MS-222, Sigma-Aldrich). Flow of red blood cells through the vasculature was confirmed before beginning the dissection as an indicator of good health. A portion of agarose was removed to expose the dorsum from the posterior hindbrain to the anterior optic tectum. The dissection was initiated by puncturing the thin epithelial layer over the rhombencephalic ventricle above the hindbrain^48^ with a sharpened tungsten needle. Small incremental anterior-directed incisions were then made along the midline as close to the surface as possible until the brain was exposed from the hindbrain entry to the middle of the optic tectum (Extended Data Fig. 1). The majority of damage associated with this dissection was restricted to medial tectal proliferation zone progenitor cells^49^ that are unlikely to have integrated into functional neuronal circuits.

Dissections lasted 1–2min, upon which time the complete dish was immersed in 2.0% formaldehyde and 2.5% glutaraldehyde fixative solution (Electron Microscopy Sciences) overnight at room temperature. Following washes, larval zebrafish were cut out from the dish in a block of agarose with a scalpel and moved to a round-bottom microcentrifuge tube. Specimens were then incubated in post-fixation solution containing 1% osmium tetroxide and 1.5% potassium ferricyanide for 2hr, washed with water, washed with 0.05M maleate buffer (pH 5.15), and stained with 1% uranyl acetate in maleate buffer overnight. During the subsequent wash step with maleate buffer, larval zebrafish were freed from the surrounding agarose block and moved to a new microcentrifuge tube. Next, specimens were washed with water, dehydrated with serial dilutions of acetonitrile in water (25%, 50%, 70%, 70%, 80%, 90%, 95%, 100%, 100%, 100%) for 10 minutes each, and infiltrated with serial dilutions of a diepoxyoctane-based low viscosity resin^50^ in acetonitrile (25%, 50%, 75%, 100%) for 1hr each. The samples were then embedded in the same resin and hardened for 2–3d at 60°C. Additional solution, washing, and timing details were described previously in a step-by-step protocol^51^.

### Serial sectioning

Sections were continuously cut with a diamond knife affixed to an ultramicrotome (EM UC6, Leica) and collected with a customised automated tape-collecting ultramicrotome^35^ onto 8mm-wide and 50–75μm-thick tape (Kapton® polyimide film, DuPont). Sectioning restarts were occasionally required for three reasons: fine-tuning of tape positioning or settings is necessary at the beginning of a run; the ultramicrotome design is constrained by a cutting window depth range of ∼200μm; and diamond knives must be shifted after cutting several thousand sections to expose the sample to a fresh cutting edge before dulling impairs sectioning quality. When necessary, restarts were completed as quickly as possible (typically 1–2min) in order to minimize possible thermal, electrostatic, or other fluctuations. For the same reason, tape reels were fed continuously rather than being reloaded. This required modifying the automated tape-collecting ultramicrotome to extend the device’s main mounting plate and enlarge tape-carrying reels to accommodate the entire length needed (compare Extended Data Fig. 2a with Fig. 1E from ref. ^35^). This combination of fast restarts and continuous tape feeding was successful at maintaining a steady state across restarts.

We sectioned two larval zebrafish specimens and these represent the only two samples we have attempted to cut with the surrounding support tissue approach. The primary focus of this study was a 5.5dpf larval zebrafish sectioned with a 45° ultra diamond knife (Diatome) and a nominal sectioning thickness that averaged 60nm with a variable setting ranging from 50–70nm depending on sectioning consistency. Restarts for this sample occurred after sections 276, 3669, 6967, 10346, 12523, 12916, and 15956. Knife shifts occurred after sections 6967 and 12916. After sectioning, the tape was cut into segments with a razor blade between collected sections and adhered with double-sided conductive carbon adhesive tape (Ted Pella) to 4in-diameter silicon wafers (University Wafer), which served as an imaging substrate. In total, 17963 × ∼60nm-thick sections were spread across 80 wafers for this specimen. A thin layer of carbon was deposited onto each wafer to prevent charging during scanning EM imaging.

One potential limitation of the 5.5dpf larval zebrafish section series presented herein is the section thickness, as minimizing section thickness is an important factor in the success of axon and dendrite reconstructions^1^. Small neuronal processes (on the order of the section thickness) are difficult to reconstruct in thicker sections especially when they are running roughly parallel to the plane of the section. To be sure that our approach was not fundamentally limited to thicker sections, we sectioned the second sample—a 7dpf larval zebrafish—with a nominal sectioning thickness that remained constant at 50nm throughout the entire cutting session using a 45° histo diamond knife (Diatome). Restarts for this sample occurred after sections 296, 312, 4114, 8233, and 12333. Knife shifts occurred after sections 4114 and 12333. In total, we obtained 15046 × ∼50nm-thick sections from 15052 attempted (Extended Data Fig. 5) and spread them across 70 wafers. The thinner sections did not cause more lost material: this series contained 6 losses (0.04%; Extended Data Fig. 5d top), 25 partial sections (0.17%; Extended Data Fig. 5d middle), no adjacent losses, and 6 adjacent lost-partial or partial-partial events (0.04%; Extended Data Fig. 5d bottom).

A nominal section thickness of ∼60nm made it possible to span the entire 5.5dpf larval zebrafish brain in ∼18000 sections, as determined by finding the location of the spinal cord-hindbrain boundary^52^. The 7dpf sample was sectioned at 50nm, but was not made the focus of imaging because of an over-trimming error that caused less of the brain to be captured. However, improved reliability for this sample despite a ∼17% reduction in nominal sectioning thickness suggests that higher axial resolution is readily attainable, though expanding the tape-carrying capacity of the microtome further may be necessary. A section thickness of ≤30nm would increase confidence in the ability to reconstruct complete neuronal circuit connectivity and this thickness will be the target in our future work. Thicknesses of ≤30nm are possible for mammalian brain sections of similar areas^36,53^, suggesting that the approach described herein will not be a limiting factor.

Once wafers contained tape segments, they were made hydrophilic by glow discharging very briefly, post-section stained for 1–2min inside a chamber containing sodium hydroxide pellets with a stabilised lead citrate solution (Leica, UltroStain II) filtered through a 0.2μm syringe filter, and then washed thoroughly with boiled water.

### Electron microscopy

WaferMapper software was used with light-based wafer overview images to semi-automatically map the positions of all sections and relate them to per-wafer fiducial markers that enabled targeted section overview acquisition at a resolution of 758.8×758.8×60nm^3^vx^−1^. Semi-automated alignment of section overviews in WaferMapper then permitted targeting for higher resolution imaging^35^.

Field emission scanning EM of back-scattered electrons was primarily conducted on a Zeiss Merlin equipped with a large-area imaging scan generator (Fibics) and stock back-scattered electron detector. An accelerating voltage of 5.0kV and beam current of 7–10nA was used for most acquisition. Imaging of back-scattered electrons at the highest resolutions (4×4×60nm^3^vx^−1^) was performed on an FEI Magellan XHR 400L with an accelerating voltage of 5.0kV and a beam current of 1.6–3.2A. Fields of view acquired on a given section varied depending on the area occupied by the sample in cross-section. All acquisition was performed with a scan rate at or under 1Mpx s^−1^. This resulted in acquisition times of 5.4 days for section overview images (758.8×758.8×60nm^3^vx^−1^), 97 days for full transverse images (56.4×56.4×60nm^3^vx^−1^), and 100 days for high-resolution brain atlas images (18.8×18.8×60nm^3^vx^−1^).

### Image alignment and intensity normalization

Producing anatomically consistent registration over our relatively long section series required control of region-of-interest drift, over-fitting, magnification changes, and intensities. In order to quickly assess the quality of the dataset and begin reconstructions, we initially performed affine intra- and inter-section EM image registrations with Fiji^54^ TrakEM2 alignment plug-ins^55^. These results revealed that additional nonlinear registration was required in order to compensate for distortions that were likely caused by section compression during cutting and minor sample charging during imaging. While the state-of-the-art elastic registration method^56^ also provided in Fiji^54^ as a TrakEM2 alignment plug-in was capable of achieving excellent local registration, we experienced difficulty—at least without modification to the existing implementation—in achieving an anatomically consistent result that preserved the overall structure of the larval zebrafish, largely due to our inability to successfully constrain region-of-interest drift and magnification changes. Furthermore, we determined that the similar AlignTK^11^ method, which uses Pearson correlation as the matching criterion coupled with spring mesh relaxation to stabilize the global volume, was likely to suffer from similar problems and would require substantial additional data handling to operate on our multi-resolution dataset.

In order to preserve the overall structure of the larval zebrafish and simultaneously achieve high-quality local registration, we turned to a new Signal Whitening Fourier Transform Image Registration (SWiFT-IR) method^53,57^. Compared to conventional Pearson or phase correlation-based registration approaches, SWiFT-IR produces more robust image matchings by using modulated Fourier transform amplitudes, adjusting its spatial frequency response during the image matching steps to maximize a signal-to-noise measure that serves as its main indicator of alignment quality. This more reliable alignment signal in turn better handles variations in biological content and typical data distortions. Furthermore, SWiFT-IR achieves higher precision in block matching as a result of the signal whitening, improves computational speeds thanks to computational complexity advantages of the fast-Fourier transform, and reduces iterative convergence from thousands to dozens of steps. Together, these capabilities enable a model-driven alignment in place of the usual approach of comparing and aligning a given section to a pre-selected number of adjacent sections.

The SWiFT-IR model we used consisted of an estimate of local aligned volume content formed by a windowed average, typically spanning ∼6 μm along the z axis orthogonal to the sectioning plane. Any severely damaged regions, in particular partial sections, were removed from the model so that they would not adversely influence alignment results. This model then served as a template during registration, where raw images were matched to the current model rather than directly to a subset of nearby sections. Alignment proceeded in an iterative fashion starting at the lowest resolution acquired (section overviews, 758.8×758.8×60 nm^3^vx^−1^) and progressing incrementally to the highest resolution image set (56.4×56.4×60 nm^3^vx^−1^ for most regions outside the brain, 18.8×18.8×60 nm^3^vx^−1^ for most regions inside the brain).

At each resolution, the source images were iteratively aligned to the current model until no further significant gain in alignment could be achieved, as indicated by the SWiFT-IR signal-to-noise figure-of-merit. The model was then transferred to the next higher resolution data by applying the current warpings to source data for that scale. Iterative model refinement then continued at this subsequent resolution level. Although the majority of computations were locally affine, residual nonlinear deformations, particularly at the highest resolutions, were represented by a triangulation mesh that deformably mapped raw data onto the model volume.

Importantly, access to the lowest resolution section overview data for each section permitted us to build an initial model that constrained subsequent registration steps to the overall structure of the larval zebrafish. Although the resolution and signal quality of overview images were intentionally kept low in favour of rapid acquisition, the fact that overviews were all quickly captured with the same microscope settings and included surrounding support tissue provided key constraints for model refinement that resulted in a more accurate global result.

More specifically, the full 17963 section overview image set was processed using SWiFT-IR to produce an initial aligned overview model at 564×564×600 nm^3^vx^−1^ resolution. Although the lowest resolution section overview images were each captured at 758.8×758.8×60 nm^3^vx^−1^, the relative oversampling in the stack direction enabled a geometrically accurate model at 564×564×600 nm^3^vx^−1^ resolution. This initial model was then cropped to the bounds of the larval zebrafish and warped, using SWiFT-IR–driven matching across the midline axis, to remove cutting compression, rotations, and other systematic variations in the pose of the specimen. The 16000 section 56.4×56.4×60 nm^3^vx^−1^ image set was next downsampled to 564×564×600 nm^3^vx^−1^ to match the overview model. It was then aligned to the overview model, resulting in an improved model. The matching and remodeling process was iterated at this scale until there was no further improvement in the SWiFT-IR match quality. The final model at this scale was then expanded to 282×282×300 nm^3^vx^−1^, or one-sixth scale, and similarly aligned in an iterative fashion until there was no further improvement. This one-sixth scale model volume (∼6 Gvx, 1600×1400×2667 vx) was convenient for rapid viewing to identify and manually correct defects and refine the pose. Further scales at 169.2×169.2×180 nm^3^vx^−1^ and 56.4×56.4×60 nm^3^vx^−1^ were similarly processed by successively expanding the model and aligning at each resolution until there was no further significant improvement in the figure-of-merit. Finally, the 12546-section highest resolution 18.8×18.8×60nm^3^vx^−1^ image set was similarly registered using the final 56.4×56.4×60nm^3^vx^−1^ volume as its model.

Image intensity was adjusted across sections for each dataset to achieve a consistent background level chosen as the average over a tissue-free region, typically a 256×256px^2^ area above and to the right of the larval zebrafish. Many images were acquired at 16-bit depth and were converted in this process to 8-bit depth. The target background level was mapped to intensity 250, which left headroom for bright pixels while keeping most tissue of interest from saturating the range. Next, a linear intensity fit between the background and a second level, typically the average grey level of a variable size continuous trajectory region on the right side of the brain, was made to adjust the intensity values for each section.

More information on SWiFT-IR software tools is publicly available online (http://www.mmbios.org/swift-ir-home).

### Image annotation and neuron reconstructions

Reconstructing neuronal structures across multi-resolution ssEM image volumes acquired from the same specimen profits from being able to simultaneously access and view separate but co-registered datasets. Without this ability, some of the time benefits of our imaging approach would be offset by the need to register and track structures across volumes that span both low-resolution, large fields of view and high-resolution, specific regions of interest. With this in mind, we added a feature to the Collaborative Annotation Toolkit for Massive Amounts of Image Data (CATMAID) neuronal circuit mapping software^58,59^ to overlay and combine image stacks acquired with varying resolutions in a single viewer (Extended Data Fig. 6). Multiple image stacks are rendered using WebGL, which makes it possible to present co-registered stacks of different resolutions in the same view. Furthermore, this feature combines stacks via a configurable overlay order, introduces blending operations for each overlaid stack, and enables programmatic shaders for dynamic image processing. When resolutions of overlaid stacks differ, the nearest available zoom level for each stack is interpolated either bilinearly or via nearest neighbours. Missing regions in each stack can be either omitted or rendered with nearest neighbour interpolation. To account for the increase in data storage and bandwidth when viewing multiple image stacks, the CATMAID image data hierarchy was extended with a shared graphics card memory cache of image tiles using a least-recently-used replacement policy. All additions and modifications to the CATMAID software are now incorporated into the main release version (http://github.com/catmaid/CATMAID).

Manual reconstruction was conducted using our modified version of CATMAID by placing nodes near the centre of each neuronal structure on every section in which it could be clearly identified. This led to a wire-frame model of each annotated structure. Starting points for reconstruction (“seeds”) of myelinated processes were manually identified by searching all tissue on a given section from top-left to bottom-right at the highest available resolution for profiles surrounded by the characteristic thick, densely stained outline associated with staining of the myelin sheath^60^ (see Fig. 1e,g,i). To obtain seeds for reconstruction of the complete projectome for one larval zebrafish, this process was repeated every 50 sections throughout the set of 16000 imaged at 56.4×56.4×60nm^3^vx^−1^ resolution. Many such reconstructions were produced in an initial, affine-only image alignment space before being mapped into the final, nonlinear image alignment space. The reconstructions reported here represent ∼450 days of human annotation.

For visualization and reported length measurements, each mapped wire-frame was smoothed using custom python-based software that implemented a Kalman smoothing algorithm on a space defined by manually annotated points within unique segments. Initial state variables for smoothing were derived by an optimization of point-to-line distance to connected reconstructed segments. Other variables were tuned with the Estimation Maximization algorithm of the pykalman library to compensate for a lack of human input where data was unavailable due to lost or partial sections. While the final image alignment was high quality, smoothing the reconstructions in this manner should produce a slight underestimate rather than an overestimate of reported reconstruction path lengths.

Neurons with known projection patterns or identities were named accordingly in the CATMAID database. For example, the reconstruction of a neuron innervating the right anterior macula (utricle) was named as “Ear_AnteriorMacula_R_01”, while an identified neuron such as the left Mauthner neuron was named “Mauthner_L”. Two identifiable neurons on each side belonged to the “MeM” class, which emanates from the nucleus of the medial longitudinal fasciculus. On each side, these were differentiated into dorsal (MeMd) and ventral (MeMv) subclasses based on consistent soma positioning.

### Correspondence across light and EM datasets

Correspondence of individual neurons or, in the case of the Z-Brain atlas, small regions across imaging modalities was achieved with landmark-based three-dimensional (3D) thin-plate spline warping of each fluorescence dataset to the ssEM dataset using BigWarp^61^. For matching *in vivo* two-photon laser-scanning microscopy data to ssEM data from the same specimen, we primarily chose landmarks consisting of distinctive arrangements of low-fluorescence regions where GCaMP5G was excluded and could be easily matched to similar patterns of neuronal nuclei at the EM level. For matching the Z-Brain atlas^37^ to the ssEM dataset, we chose landmarks based on identifiable structures—such the region boundaries, known clusters of neurons, midline points, ganglia, and the outline of the brain—in the Z-Brain averaged *elavl3*:H2B-RFP or anti-tERK fluorescence image stacks that were also observed in the ssEM data. The same Z-Brain landmarks were used for transforming a version^62^ of the Zebrafish Brain Browser^38^ that was previously registered into the Z-Brain atlas.

### Data and reconstruction visualizations

The image volumes and reconstructions were visualized using Vivaldi^63^, a domain-specific language for rendering and processing on distributed systems, because it provides access to the parallel computing power of multi-GPU systems with language syntax similar to python.

For volume visualizations, we used a direct volume rendering ray-casting technique in which an orthogonal or perspective iterator was marched along a viewing ray while sampled voxel colours were accumulated using an alpha compositing algorithm. We screened out regions containing only support tissue during rendering with labelled volumes constructed by interpolating between manually produced masks indicating which image voxels belong to each separate tissue region. In cases where separate image volumes of the same region were rendered together (such as when ssEM data and fluorescence data were combined), direct volume rendering was performed by combining front-to-back colour and alpha compositions formed from the different transfer function belonging to each image volume.

For volume visualizations that included reconstructions, direct volume rendering of image data was combined with streamline rendering of reconstructions using two different techniques. The first combined an OpenGL framebuffer with the Vivaldi volume rendering. In this case, each streamline was rendered using OpenGL as a tube into an off-screen buffer (i.e., Framebuffer Object). Vivaldi then compared the resulting render and depth buffers to perform direct volume rendering of only the image data above the streamline depth value. This made it possible to ignore any image data voxels located behind streamlines, which were treated as opaque. The second technique involved generating a complete streamline volume by 3D rasterization. This streamline volume was then combined with the image data volume for direct volume rendering. The former technique is faster and can cope with dynamic streamline changes, but the latter was found to yield better overall rendering quality. Visualizations of reconstructions without the image volume context were similarly rendered as streamlines in Vivaldi or in the CATMAID 3D WebGL viewer.

Reference plane (e.g., horizontal, sagittal, and section) images were rendered by detecting the zero-crossing of each viewing ray and the plane. Support for viewing these opaque data views in some spatial regions alongside the semi-transparent volume visualization views in other regions was introduced as a new function (clipping_plane) in Vivaldi. Similarly, contour (nonplanar) reslice support was developed to illustrate a flattened view along a specific contour (or neuron path) consisting of vertical line segments extracted from image data.

For many cases, the size of the image volume being rendered was larger than the available memory. In order to support out-of-core processing, we developed and integrated into Vivaldi a slice-based streaming computing framework using Hadoop distributed file system (HDFS) that will be reported elsewhere.

### Symmetry analyses

Initial observations of apparent symmetry in subsets of myelinated axon reconstructions were found during visual inspection (Fig. 3; Supplementary Video 8; Supplementary Video 9). To more quantitatively assess the extent of symmetry, we developed methods for 3D symmetry plane fitting and two symmetry analyses: one that produces a cost associated with the shape and position similarity between reconstructed 3D structures and another that compares the relative two-dimensional (2D) positioning of identified reconstruction pairs on one side with the relative positioning for the same identified pair on the opposite side at a given cross-section level (Fig. 4d). Only the longest reconstructed path from the soma through the myelinated axon projection was considered in plane fitting and subsequent analyses. Reconstructed dendrites or short axonal branches were ignored. Each resulting reconstruction path (skeleton) was represented as an ordered list of nodes (points) taken directly from manual reconstructions. Sidedness (left/right) for each reconstruction was determined by its soma position.

Most of these new analyses approaches, including symmetry plane fitting and 3D symmetry comparisons, are described elsewhere^64^. The symmetry plane fitting, in brief, involves choosing an approximate symmetry plane, reflecting the complete set of points belonging to the reconstruction subset of interest with respect to this plane, registering the original and reflected point clouds with an iterative closest point algorithm, and inferring the optimal symmetry plane from the reflection and registration mappings. The subset of reconstructions from which this plane fitting was performed consisted of all identified neurons whose axon projections formed part of the medial longitudinal fasciculus (MLF), determined primarily using refs. ^40,41^.

The 3D symmetry comparison for each given template reconstruction on one side, in brief, involves reflecting all reconstructions from the opposite side and computing a matching cost via dynamic time warping (DTW) for each comparison. For our purposes, the DTW cost is taken as the sum of the Euclidian distances between all matched points normalized to the number of matched point pairs (Extended Data Fig. 9a-c). The DTW gap cost parameter for matching a point in one sequence with a gap in another is set to zero because our data is sampled at a nearly constant rate and we seek the optimal subsequence match even in cases where one sequence is shorter than or offset with respect to the other. To compensate for unmatched regions (i.e., overhangs), the DTW cost is then multiplied by a penalty factor that is proportional to the lengths of the sequences that remain unmatched (total length divided by matched length). Comparing each reconstruction on one side to all reconstructions from the opposite side formed a cost matrix from which an optimal pairwise assignment could be determined without any bias introduced from the previously determined identities. The Munkres algorithm^65^ (also known as the Hungrarian method) was used to compute the globally optimal pairwise assignment between reconstructions. For visualization clarity and to decrease analysis time, the subset of reconstructions to which this 3D symmetry analysis was applied was restricted to identified neuron classes with only 1-2 members per side whose axon projections formed part of the MLF, determined primarily using ref. ^41^.

We next sought to compare the relative 2D positioning of axon pairs with the relative 2D positioning of the same pair on the opposite side at a given cross-section level (Fig. 4d). For visualization clarity, the subset of reconstructions to which this relative 2D positioning analysis was applied was restricted to the Mauthner cell and the identified neurons whose somata resided in the nucleus of the medial longitudinal fasciculus.

First, a small angle offset in the sectioning plane relative to the true transverse plane was compensated for by projecting the point coordinates of reconstructions such that the previously computed symmetry plane became the plane *x* = 0. Given the transverse planes *z*_0_, *z*_1_, and a projected *skeleton S* containing points *s* = (*s*_*x*_, *s*_*y*_, *s*_*z*_), we let 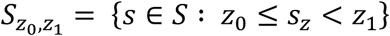. That is, 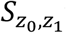 was taken as the subset of points from *S* whose coordinates *s*_*z*_ are all in the interval [*z*_0_, *z*_1_). We refer to the subset of ℝ^3^ bounded by *z* ∈ [*z*_0_, *z*_1_) as the *slice* [*z*_0_, *z*_1_). For each slice [*z*_0_, *z*_1_) and skeleton *S*, we defined 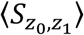 as the mean of the elements in 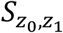. This mean was then taken as representative of the skeleton *S* in slice [*z*_0_, *z*_1_) both for further analysis and plotting. Note that all analysis and plotting presented in static form (Fig. 4) was based on a slice thickness corresponding to a single section (∼60nm), in which case each slice consisted simply of each pair of adjacent sections. This was not the case for dynamic presentation (Supplementary Video 10) due to video time and size constraints.

For comparing one pair with its counterpart (Extended Data Fig. 9e), we then took *s*_1_, …, *s*_*n*_ to be the set of representative points in a fixed slice for skeletons *S*_1_, …, *S*_*n*_ and took *t*_1_, …, *t*_*n*_ to be the representative points (for the same slice) of the respective skeletons *T*_1_, …, *T*_*n*_ that were previously paired to *S*_1_, …, *S*_*n*_ by the Munkres algorithm assignment in the 3D symmetry analysis, which for this subset of reconstructions always implied assignment of each identified neuron to its homolog on the opposite side.

To quantify the degree of similarity between any given pair and its opposite side counterpart, we devised two measures (Extended Data Fig. 9e). The first, termed the *angle difference*, *a*_*i,j*_, between pairs, is defined as:

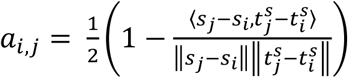

The second, termed the *distance difference*, *d*_*i,j*_, between pairs, is defined as:

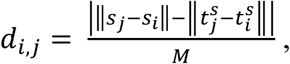

where *i*, *j* are skeleton indices, 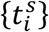 is the reflections of {*t*_*i*_} with respect to the computed plane of symmetry, and *M* is the maximum of 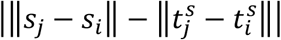 across all pairs and all stacks. Note that *a*_*i,j*_ and *d*_*i,j*_ are normalized such that they can vary from 0 (no difference, 0° or 0μm) to 1 (maximum difference, 180° or ∼8μm). Further, when the points *s*_*i*_ and *s*_*j*_ are perfectly symmetric with respect to points *t*_*i*_ and *t*_*j*_, then *a*_*i,j*_ = 0 and *d*_*i,j*_ = 0.

To more easily visualize this quantification, a *difference matrix*, *D*, was generated for each slice such that *D*(*i*,*j*) = *a*_*i,j*_ if *j* > *i* and *D*(*i*,*j*) = *d*_*i,j*_ if *j* < *i* (Fig. 4e; Extended Data Fig. 9f; Supplementary Video 10). Calculating the variance for each element in *D* across all slices showed which pairs deviated most with respect to the opposite side counterpart (Fig. 4j). Heatmaps of vectorised (linearized) upper (*j* > *i*) and lower (*j* < *i*) triangles of *D* across slices with the same pair ordering additionally revealed where differences were present for single pairs and their opposite side counterparts (Fig. 4g-h; Extended Data Fig. 9g-h; Supplementary Video 10). Plotting the sum of all *a*_*i,j*_ and *d*_*i,j*_ values for a given slice further illustrated where differences were present across the entire subset (Fig. 4i). Finally, the exact same process was performed after swapping the identity (assignment) of the two reconstructions with the lowest 3D symmetry analysis costs (MeLc and Mauthner neurons) to provide a basis for comparison (Fig. 4i).

### Online data hosting

All aligned ssEM data, reconstructions, co-registered Z-Brain data, and a guide outlining ways to interact with the data are publicly available online (http://hildebrand16.neurodata.io/). Our enhanced version of CATMAID^58,59^ software with multi-resolution support is used for visualization (Extended Data Fig. 6). All image data is served as a series of 8-bit 1024×1024px^2^ PNG images with a tRNS value of 255 specified to enable transparency. To provide a smooth visualization experience, the original resolution for each image stack was down-sampled multiple times to create a resolution hierarchy, where each level in the hierarchy corresponds to an image that is half the size as in the previous level. The entire down-sampled dataset required 2.8 TB of disk space (608 GB for 56.4×56.4×60nm^3^vx^−1^ resolution EM data, 1.8 TB for 18.8×18.8×60nm^3^vx^−1^ resolution EM data, 356 GB for 4×4×60nm^3^vx^−1^ resolution of dorsal neuromasts, and 29 GB for 300×300×300nm^3^vx^−1^ resolution of Z-Brain data). To serve data to the end user, we used the Amazon Web Services (AWS) Simple Storage Service (S3). The AWS S3 built-in web server was used to serve the static image files. A public CATMAID instance was then deployed on Amazon EC2 to point to the images in S3.

Custom tools generated for data handling and analysis are publicly available (http://github.com/davidhildebrand/hildebrand16/).

## Acknowledgements

We thank D.D. Bock, K.-H. Huang, and P. Huang for preliminary studies; L.-H. Ma and M.B. Ahrens for dissection help; E. Raviola, H.S. Kim, J.A. Buchanan, E.J. Benecchi, and S. Ito for tissue preparation guidance; K.J. Hayworth, J.L. Morgan, N. Kasthuri, and R. Schalek for sectioning and scanning electron microscopy advice; T. Kazimiers and J.A. Bogovic for software assistance; the Harvard Image and Data Analysis Core (H.L. Elliot, T. Xie, D.L. Richmond) and D. Berger for image processing assistance; the Harvard Center for Brain Science Neuroengineering Core (E.R. Soucy, J.S.F. Greenwood) for instrumentation support; and B.L. Shanny, A.M. Roberson, F. Gao, M.A. Afifi, F. Camacho Garcia, A.D. Wong, C.S. Elkhill, T.J. Cawley, R.J. Plummer, K.M. Runci, A. Haddad, P.E. Lewis, I. Odstrcil, A.H. Cohen, M.D. Petkova, and P.I. Petkova for reconstructions. This work was supported by the NIH through the NINDS to F.E. (DP1 NS082121, RC2 NS069407) and to W.-C.A.L. (R21 NS085320) and to the Harvard Center for Brain Science Neuroengineering Core (P30 NS062685), through the NIGMS to MMBioS via resources at the Pittsburgh Supercomputing Center (P41 GM103712), through the NCRR to the Orchestra High-Performance Compute Cluster (S10 RR028832); by the Defense Advanced Research Projects Agency (DARPA) SIMPLEX program through SPAWAR to R.B. and J.T.V. (contract N66001-15-C-4041); by the National Research Foundation of Korea (NRF) through the Bio & Medical Technology Development Program, MSIP (NRF-2015M3A9A7029725), and the Ministry of Education Basic Science Research Program (NRF-2014R1A1A2058773) to W.-K.J.; and by training fellowships from the NIH (T32 MH20017, T32 HL007901) and NSF (IIA EAPSI award 1317014) to D.G.C.H.

## Author Contributions

F.E. designed the study. R.P., I.H.B., O.R., and D.G.C.H. conducted light microscopy. D.G.C.H. prepared and sectioned samples. D.G.C.H. and G.S.P. performed electron microscopy. A.W.W., S.S., and D.G.C.H. aligned electron micrographs. D.G.C.H. and O.R. completed correspondence across imaging modalities. D.G.C.H. managed neuron reconstructions. R.M.T., B.J.G., and D.G.C.H. post-processed and analysed reconstructions. M.C. and D.G.C.H. performed symmetry analyses. W.C., T.M.Q., J.M., D.G.C.H., and W.-K.J. produced visualizations. A.S.C. modified hosting and annotation software. A.B., K.L., R.B., and J.T.V. provided dataset hosting. F.E., J.W.L., W.-C.A.L., and A.F.S. managed various aspects of the project and provided resources. D.G.C.H., F.E., and J.W.L. wrote the manuscript with input from other authors.

## Author Information

The authors declare no competing financial interests. Correspondence and requests for materials should be addressed to D.G.C.H. or F.E.

**Extended Data Figure 1:**
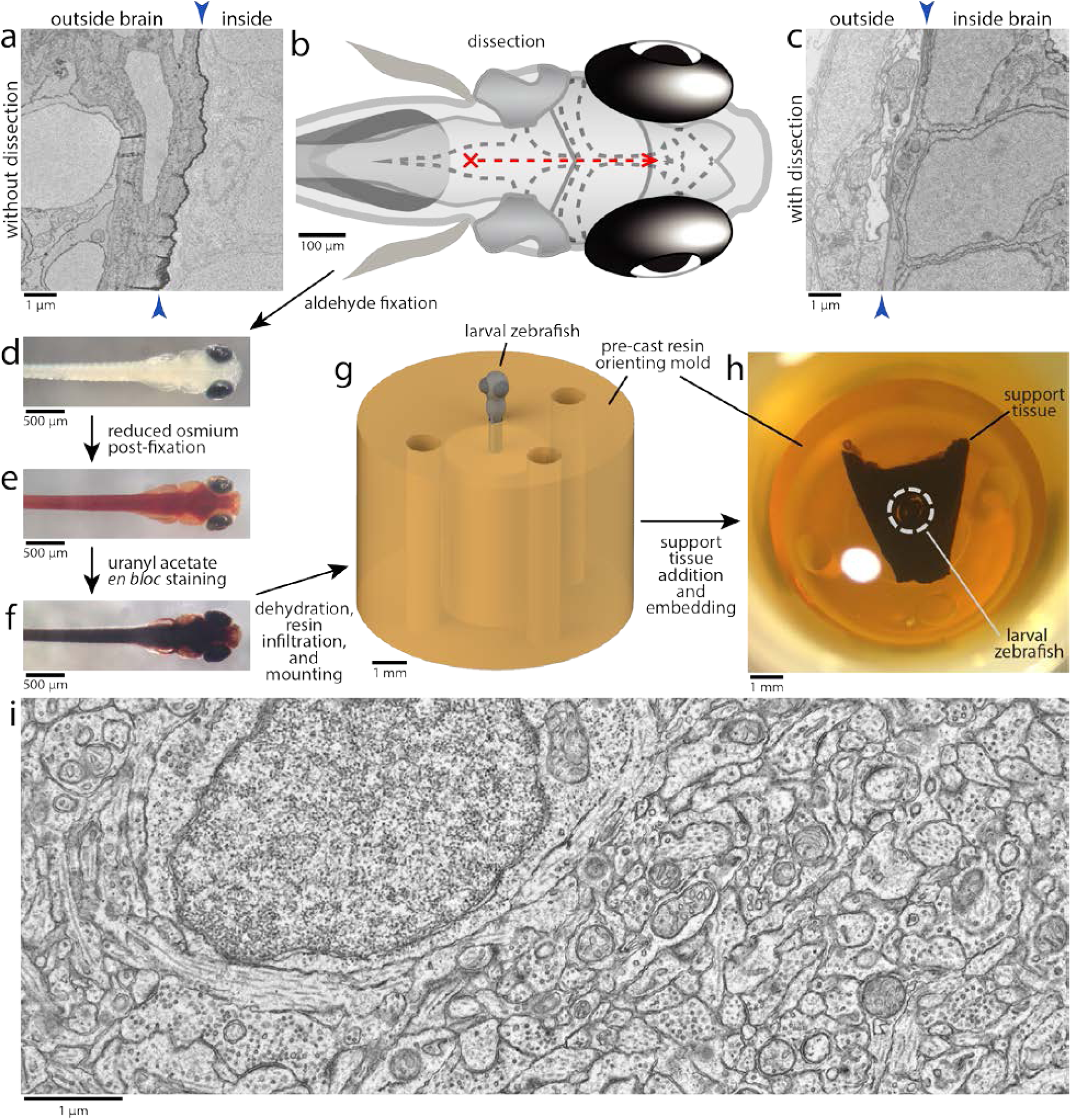
Preparing larval zebrafish brain tissue for electron microscopy. **a**, Immersion of intact specimen into tissue processing solutions typically resulted in poor preservation of brain ultrastructure due to a membrane diffusion barrier (arrowheads). **b–c**, Dissecting away skin and membranes permitted diffusion of solutions, resulting in improved preservation of brain ultrastructure. To minimize damage, dissections were initiated by puncturing the rhombencephalic ventricle dorsal to the hindbrain with a sharpened tungsten needle (red cross). Small anterior-directed incisions along the midline as close to the surface as possible were then made until the brain was exposed from the hindbrain to the anterior optic tectum (red dashed line). **d–f**, Following dissection and aldehyde fixation (**d**), samples were post-fixed with a reduced osmium solution (**e**) and stained with uranyl acetate (**f**). **g–h**, Processed larvae were then dehydrated with acetonitrile, infiltrated with a low-viscosity resin, mounted in a micromachined pre-cast resin mold to orient the sample for transverse sectioning (**g**), and surrounded by a support tissue (murine cortex) that was found to stabilize sectioning (**h**). **i**, Representative ultrastructure achieved using this process acquired as a transmission electron micrograph from a section through the optic tectum of an early dissection test specimen. Scale bars: **g**–**h**, 1mm; **d**–**f**, 500μm; **b**, 100μm; **a**,**c**,**i**, 1μm.

**Extended Data Figure 2:**
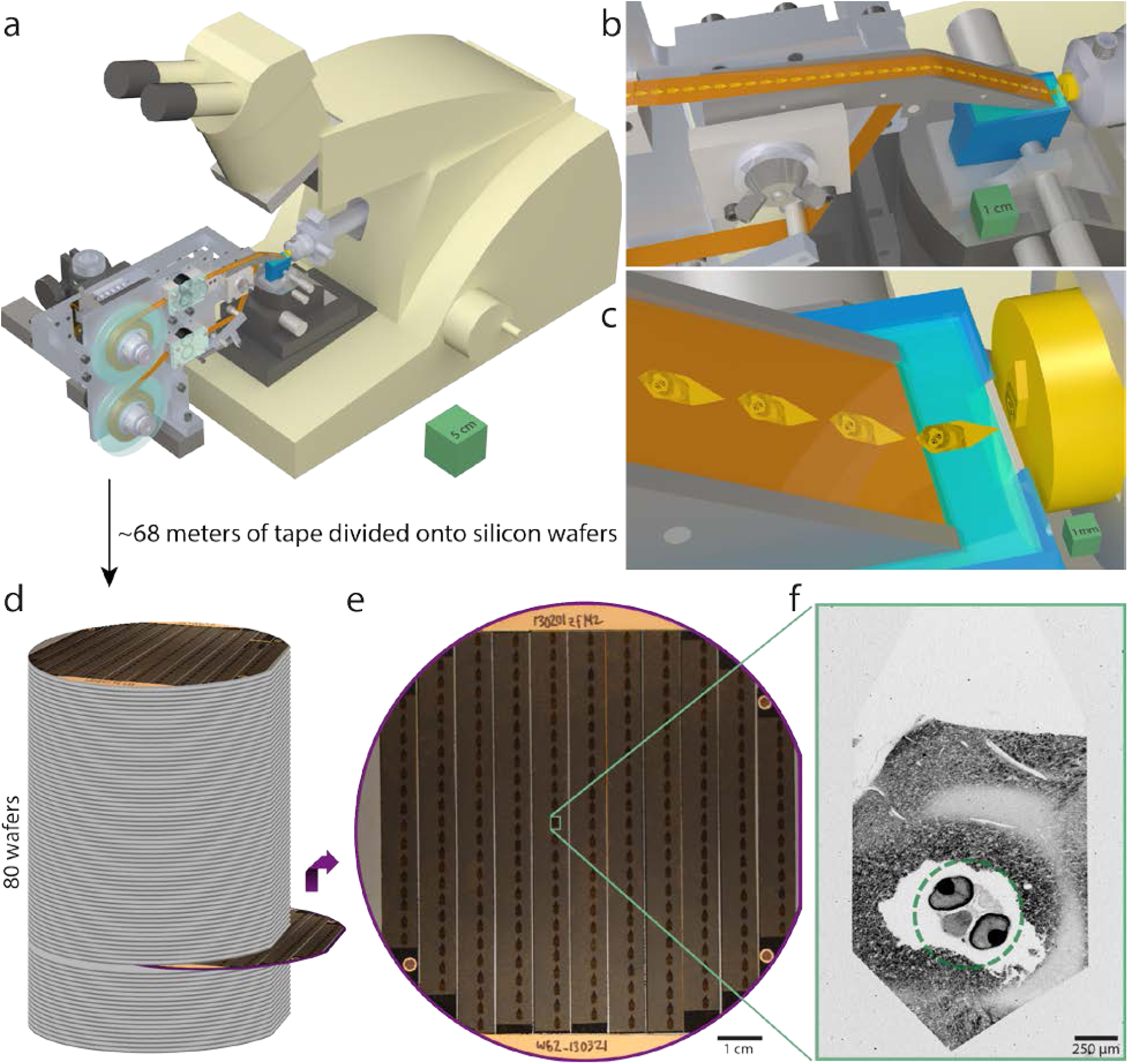
Serial sectioning and ultrathin section library assembly for a 5.5dpf larval zebrafish. **a**, Serial sections of resin-embedded samples were picked up with an automated tape-collecting ultramicrotome modified for compatibility with larger reels containing enough tape to accommodate tens of thousands of sections. **b–c**, Direct-to-tape sectioning resulted in consistent section spacing and orientation. Just as a section left the diamond knife (blue), it was caught by the tape. **d**, After serial sectioning, the tape was divided onto silicon wafers that functioned as a stage in a scanning electron microscope and formed an ultrathin section library. For a series containing all of a 5.5dpf larval zebrafish brain, ∼68m of tape was divided onto 80 wafers (with ∼227 sections per wafer). **e**, Wafer images were used as a coarse guide for targeting electron microscopic imaging. Fiducial markers (copper circles) further provided a reference for a per-wafer coordinate system, enabling storage of the position associated with each section and, thus, multiple rounds of re-imaging at varying resolutions as needed. **f**, Low-resolution overview micrographs (758.8×758.8×60nm^3^vx^−1^) were acquired for each section to ascertain sectioning reliability and determine the extents of the ultrathin section library. Scale boxes: **a**, 5×5×5cm^3^; **b**, 1×1×1cm^3^; **c**, 1×1×1mm^3^. Scale bars: **e**, 1cm; **f**, 250μm.

**Extended Data Figure 3:**
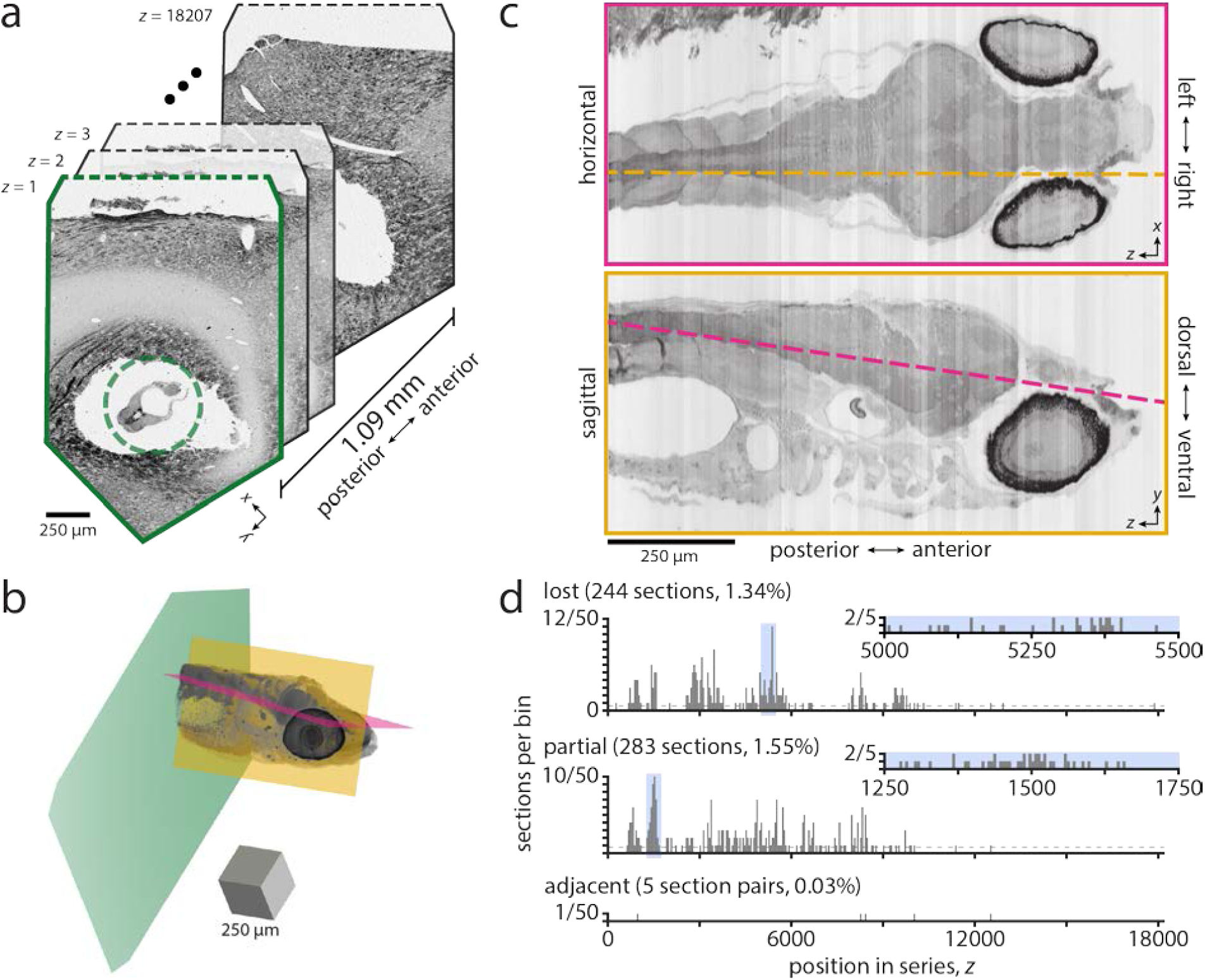
Serial sectioning through the anterior quarter of a 5.5dpf larval zebrafish. **a**, Overview micrographs (758.8×758.8×60nm^3^vx^−1^) from a collection of 18207 × ∼60 nm-thick transverse serial sections that span 1.09mm through a larval zebrafish. Embedding the larval zebrafish (green dashed circle) in a support tissue (murine cortex) stabilized sectioning. Dashed lines show where overview images were cropped. **b**, Volume rendering of aligned overview micrographs. Magenta and yellow planes correspond to reslice planes in **c**. Green plane corresponds to section outlined in **a**. **c**, Reslice planes through an aligned overview image volume reveal structures contained within the series and illustrate the sectioning plane relative to the horizontal (top) and sagittal (bottom) body planes. This series spans from myotome 7 through the anterior larval zebrafish, encompassing part of the spinal cord and the entire brain. Line colours indicate the intersection of the reslice planes. **d**, Histograms of lost, partial (missing any larval zebrafish tissue), or adjacent (lost-partial or partial-partial) events per bin of 50 sections. In total, 244 (1.34%) sections were lost and 283 (1.55%) were partial in this series. No two adjacent sections were lost. Inset histograms expand the shaded regions to provide a detailed view of sectioning reliability with bin sizes of 5 sections. Dashed lines indicate the number of lost sections if uniformly distributed throughout the series. Scale box: **b**, 250×250×250μm^3^. Scale bars: **a**,**c**, 250μm.

**Extended Data Figure 4:**
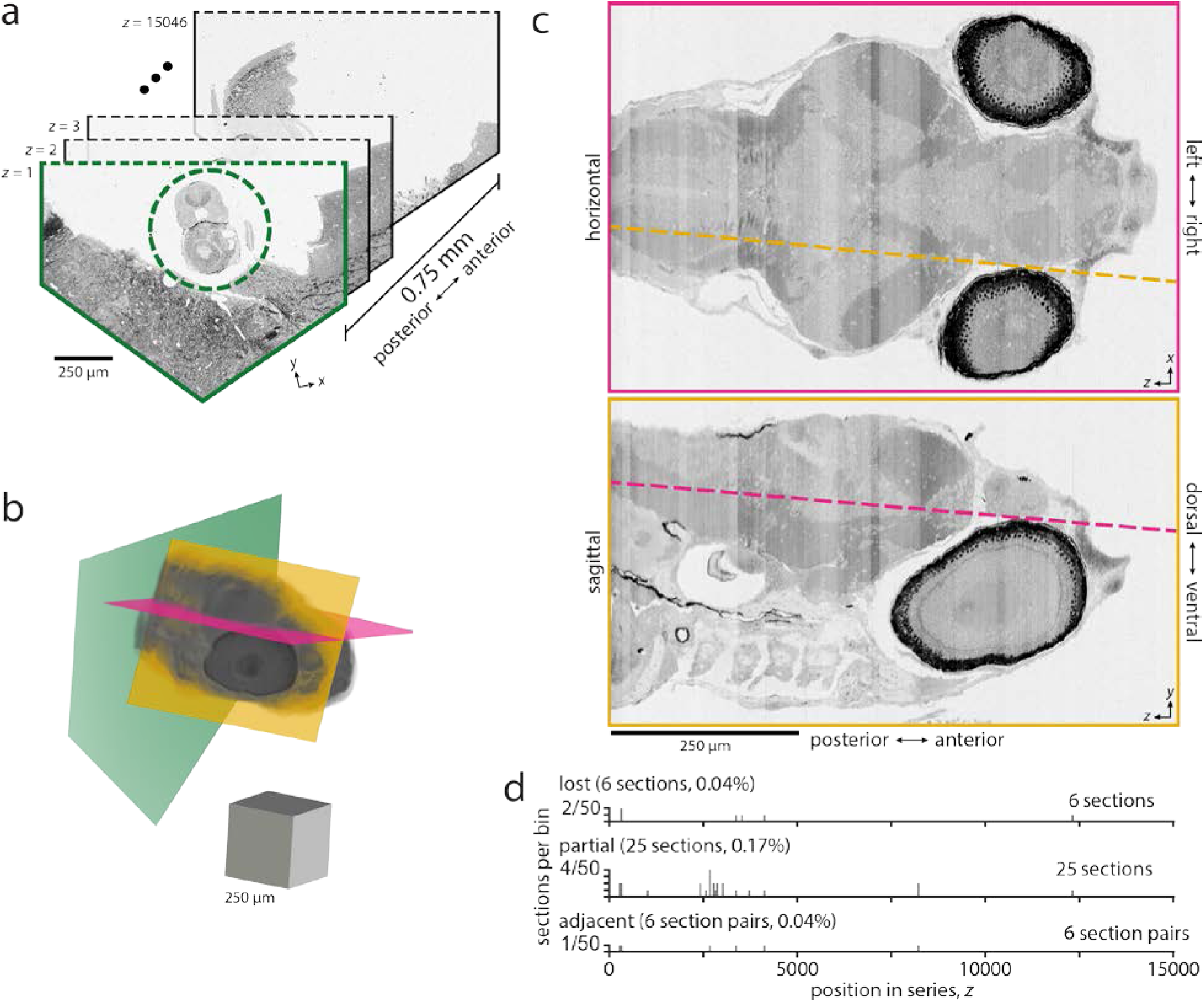
Serial sectioning through most of a 7dpf larval zebrafish brain. **a**, Overview micrographs from a collection of 15046 × ∼50 nm-thick transverse serial sections that span 0.75mm through a 7dpf larval zebrafish. Surrounding part of the larval zebrafish (green dashed circle) with a support tissue (murine cortex) stabilized sectioning. Dashed lines show where overview images were cropped. **b**, Volume rendering of aligned overview micrographs. Magenta and yellow planes correspond to reslice planes in **c**. Green plane corresponds to section outlined in **a**. **c**, Reslice planes through an aligned overview image volume reveal structures contained within the series and illustrate the sectioning plane relative to the horizontal (top) and sagittal (bottom) body planes. This series spans from posterior hindbrain through the anterior larval zebrafish, encompassing most of the brain. Line colours indicate the intersection of the reslice planes. **d**, Histograms depicting the number of lost, partial (missing any larval zebrafish tissue), or adjacent (lost-partial or partial-partial) events per bin of 50 sections throughout the series. In total, 6 (0.04%) sections were lost and 25 (0.17%) were partial in this series. No two adjacent sections were lost. Scale box: **b**, 250×250×250μm^3^. Scale bars: **a**,**c**, 250μm.

**Extended Data Figure 5:**
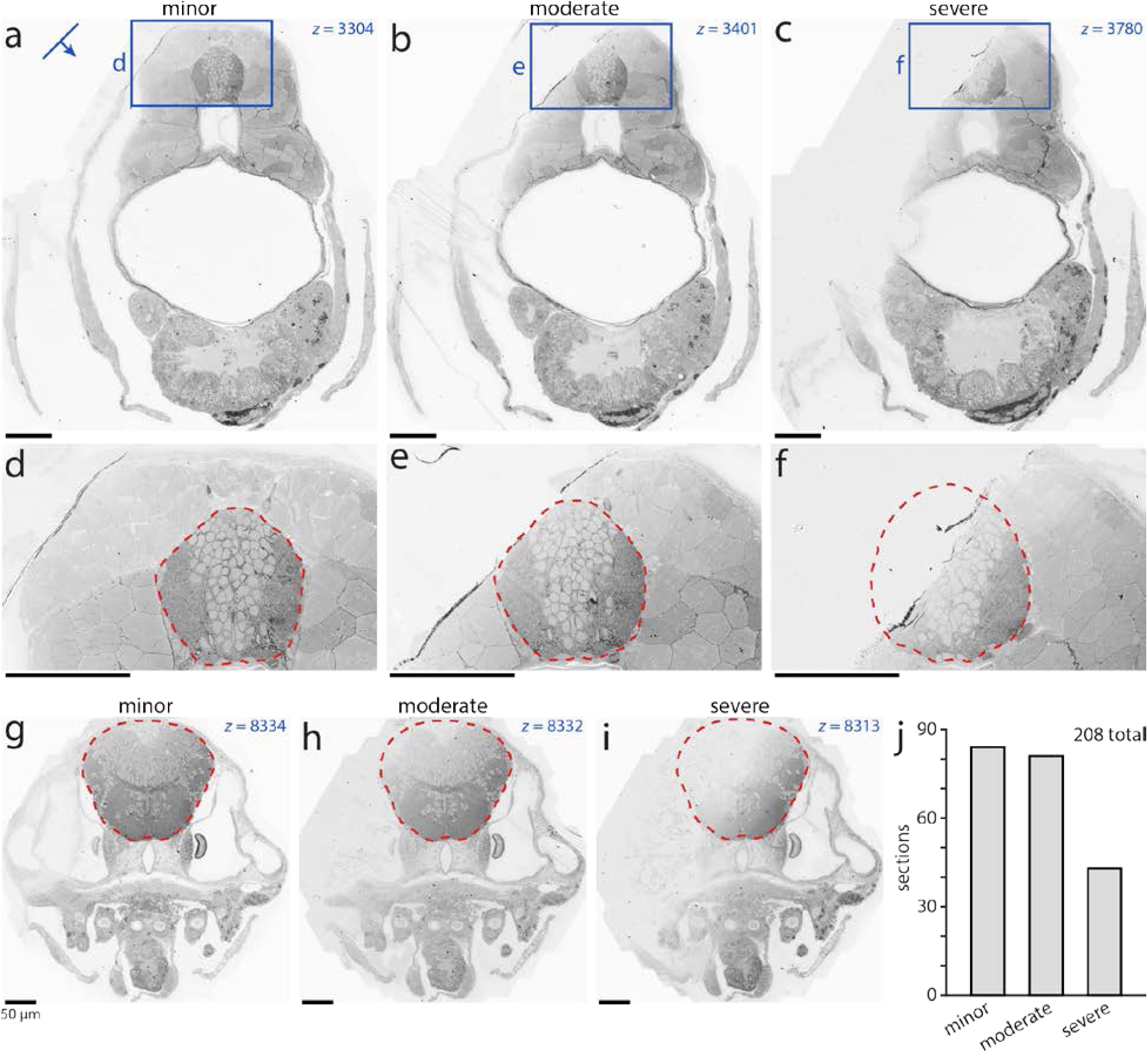
Description of partial sections. Collected sections were deemed partial if any larval zebrafish tissue appeared to be missing. In total, 283 sections out of 18207 were classified as partial. Those that were imaged at higher resolution were further categorized into minor, moderate, or severe subclasses. In minor cases, only tissue outside the brain was absent. Moderate cases lacked less than half of the brain. Severe cases were missing more than or equal to half of the brain. Note that it is possible that apparently missing tissue is contained in a slightly thicker adjacent section, in which case it is not entirely lost and may be accessible with different imaging strategies. **a–c**, Posterior examples of partial sections from each category. The line and arrow indicate the orientation and direction of sectioning. **e–f**, Expanded views of brain tissue from the sections depicted in **a–c**. Red dashed contours define the brain outline from an adjacent section. **g–h**, Anterior examples of partial sections from each category. **j**, Number of sections in each category for the 208 partial sections contained within the 16000 imaged at nearly isotropic (56.4×56.4×60nm^3^vx^−1^) resolution. Scale bars: **a**–**i**, 50μm.

**Extended Data Figure 6:**
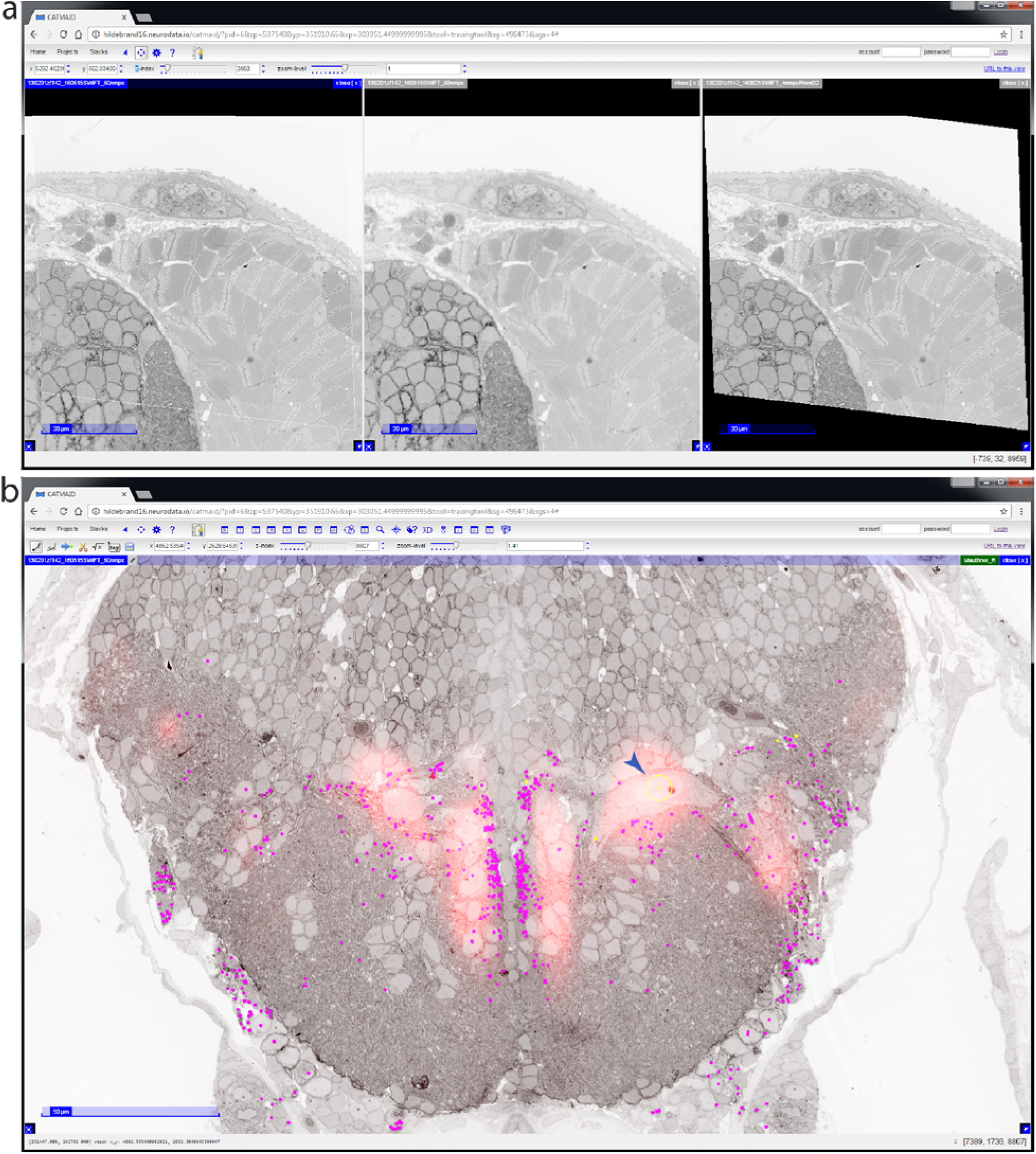
Software modifications for seamless interaction with multiple co-registered ssEM datasets and reference atlas overlays. Reconstructing neuronal structures across multi-resolution ssEM image volumes acquired from the same specimen profits from being able to simultaneously access and view separate but co-registered datasets. Without this ability, some of the time benefits of our imaging approach would be offset by the need to register and track structures across volumes that span both low-resolution, large fields of view and high-resolution, specific regions of interest. With this in mind, we added a feature to the Collaborative Annotation Toolkit for Massive Amounts of Image Data (CATMAID) neuronal circuit mapping software to overlay and combine image stacks acquired with varying resolutions in a single viewer. This feature is now available in the main CATMAID distribution. Furthermore, all datasets described here are openly accessible and hosted using this improved version of CATMAID software (http://hildebrand16.neurodata.io). **a**, Single-section images from two co-registrered ssEM datasets acquired at different resolutions from the same specimen. The combined view (left) overlays high resolution data (right, 4×4×60nm^3^vx^−1^) onto nearly isotropic resolution data (middle, 56.4×56.4×60nm^3^vx^−1^). **b**, Integrated view of multiple co-registered ssEM datasets with manual reconstructions (coloured dots) and the spinal backfill label from the Z-Brain atlas. Visible in this section is the red spinal backfill fluorescence directly on top of, for example, the Mauthner cell body (arrowhead).

**Extended Data Figure 7:**
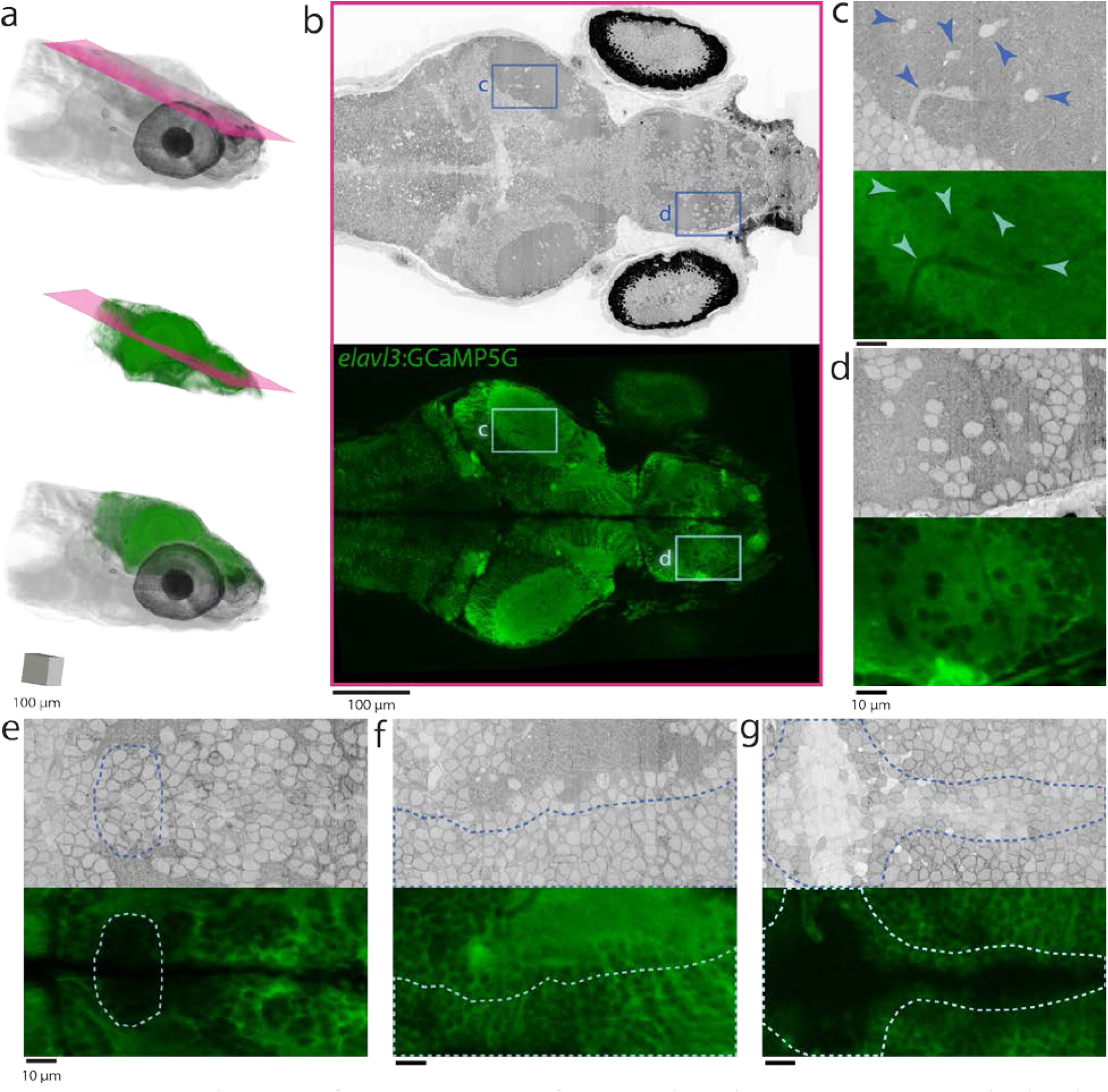
Correspondence of neuron identity across whole-brain *in vivo* light and *post hoc* ssEM datasets. Co-registration of *in vivo* light microscopy and EM datasets can be accomplished with thin-plate spline coordinate transformations guided by manually identified landmarks. **a**, Volume renderings of the ssEM dataset (top), warped *in vivo* two-photon imaging of *elavl3*:GCaMP5G fluorescence from the same specimen (middle), and a merge (bottom). Reslice planes shown in **b** are indicated by magenta planes. **b**, Near-horizontal reslice planes from the ssEM volume (top) and the warped *in vivo* light microscopy image volume (bottom) show gross correspondence throughout the brain across imaging modalities. **c–d**, Magnified views reveal single-neuron correspondence in the optic tectum (**c**) and telencephalon (**d**). **e–g**, This exercise revealed the imaging conditions, labelling density, and structural tissue features necessary for reliable matching across imaging modalities. It was difficult in regions (enclosed by dotted contours) where fluorescence signal was low (**e**), where many cells were packed closely together (**f**), and where progenitors add new neurons in between light microscopy and preparation for ssEM (**g**). Improving the light-level data with specific labelling of all nuclei and faster light-sheet or other imaging approaches should greatly improve the ease and accuracy of matching neuron identity across modalities. This ability to assign neuron identity across imaging modalities demonstrates proof-of-principle for the potential integration of rich neuronal activity maps with subsequent whole-brain structural examination of functionally characterized neurons and their networks. Arrowheads in **c** indicate the same structures as observed in each modality. Elongated structures are blood vessels. Scale box: **a**, 100×100×100μm^3^. Scale bars: **b**, 100μm; **c**–**g**, 10μm.

**Extended Data Figure 8:**
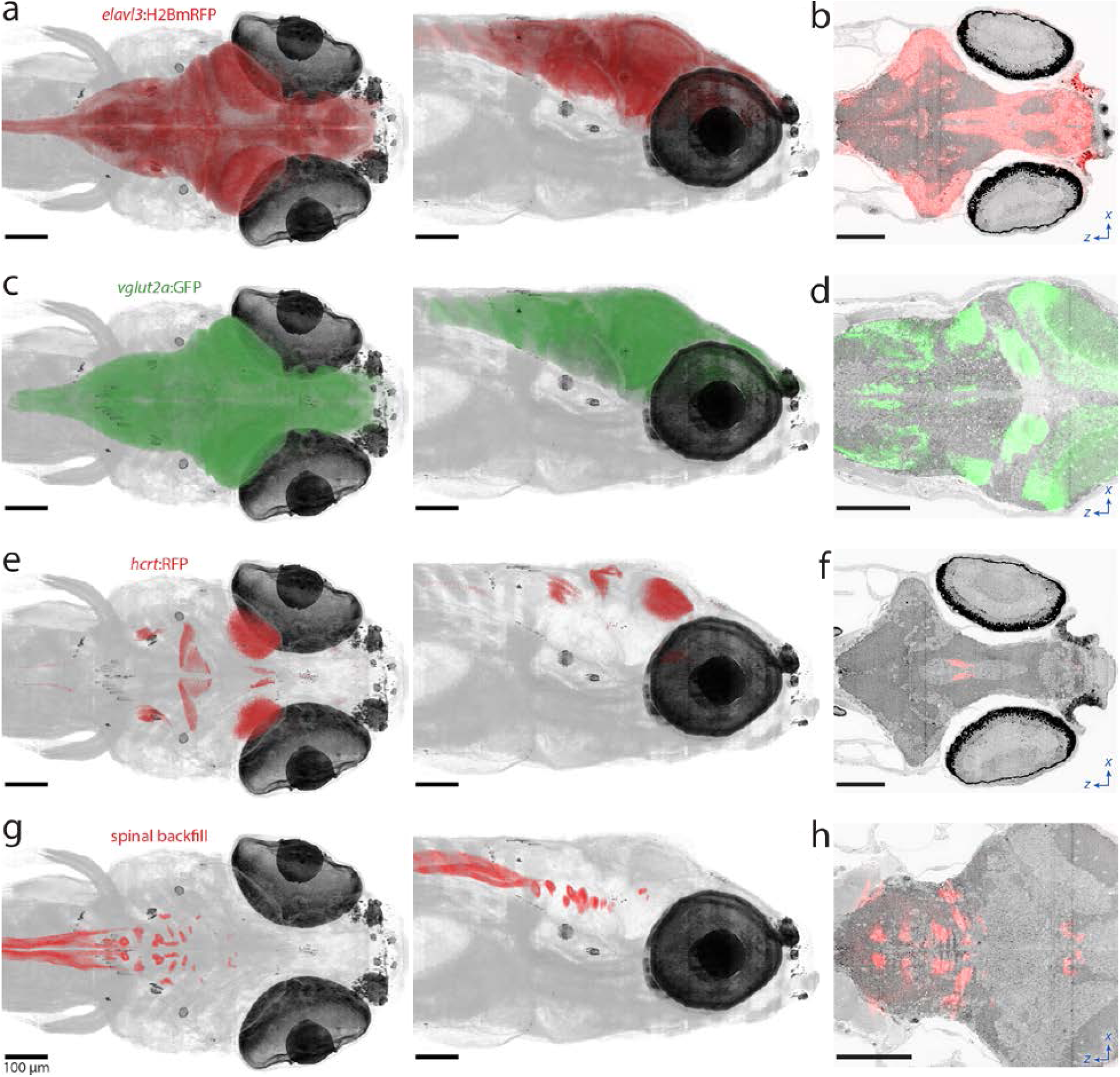
Registration of reference atlases to the multi-resolution ssEM dataset. Cross-modal registration of the Z-Brain atlas and the Zebrafish Brain Browser allows for the characterization of specific domains within the ssEM dataset defined by, for example, genetically restricted labels (**a–f**) or retrograde labeling (**g–h**). **a**,**c**,**e**,**g**, Dorsal (left) and lateral (right) views through combined, dual-volume renderings of the ssEM dataset and Z-Brain atlas image volumes of a *elavl3*:H2BmRFP transgenic line (**a**), *vglut2a*:GFP transgenic line (**c**), *hcrt*:RFP transgenic line (**e**), and spinal backfill retrograde labeling (**g**). **b**,**d**,**f**,**h**, Z-Brain atlas fluorescence signal from a *elavl3*:H2BmRFP transgenic line (**b**), *vglut2a*:GFP transgenic line (**d**), a *hcrt*:RFP transgenic line (**f**), and spinal backfill retrograde labeling (**h**) overlaid onto horizontal reslice planes through the ssEM dataset. Clearly overlapping in the spinal backfill label and ssEM reslice (**h**) are the Mauthner cell and nucleus of the medial longitudinal fasciculus neurons positions, which indicate high quality of registration. Scale bars: **a**–**h**, 100μm.

**Extended Data Figure 9:**
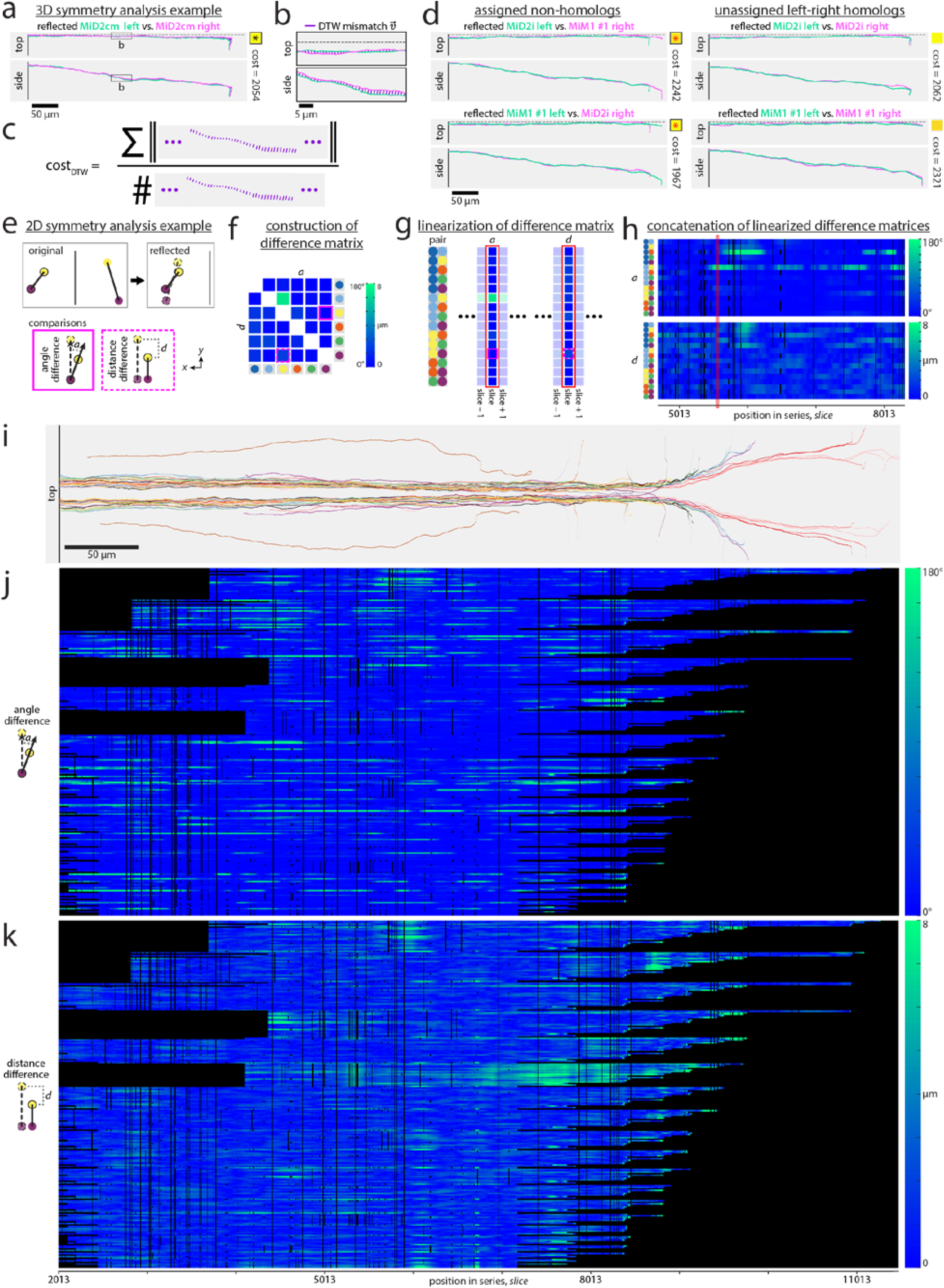
Visual description and additional examples for symmetry analysis. **a**–**c**, Three-dimensional (3D) symmetry analysis for one example pair. **a**, To compare the left MiD2cm axon to the right homolog, the left side was first reflected across the plane of symmetry (dotted line). **b**, Generating a value that represents the similarity in position and shape of the two axons was built using a dynamic time warping sequence matching approach (see Methods). **c**, Each cost was calculated as the sum of the Euclidian distances between points matched by a dynamic time warping algorithm, then normalized by the number of matches. The cost is then multiplied by a penalty factor that is proportional to the lengths of the sequences that remain unmatched (total length divided by matched length; not illustrated). **d**, One pair of myelinated axons were not assigned to their left-right homologs (Fig. 4b, red asterisks) in a globally optimal pairwise assignment for a selection of 44 identified reticulospinal neurons (22 on each side). Comparing views and costs for the assigned non-homologs (left) with unassigned left-right homologs (right) reveals similar matching costs in all cases. However, the combined cost for both non-homologs is slightly lower than that for both left-right homologs (by 174). Since the global assignment seeks to minimize the overall cost, this is likely why it assignment result grouped non-homologs over left-right homologs. The fact that the penalty accounting for the unmatched regions between the non-homologs did not increase the cost enough to prevent this assignment shows that it has a small effect in cases with short unmatched overhangs. **e**–**h**, Two-dimensional (2D) symmetry analysis of axon neighbour relations. **e**, In each 2D slice, for every pair of identified axons on one side, the same pair on opposite side was reflected across the plane of symmetry. Two metrics were then calculated to relate the original and reflected pairs: the angle difference, or dot product between the vectors connecting the pairs, and the distance difference, or the difference between the lengths of the vectors connecting the pairs. **f**, For each slice, a distance matrix was constructed from the angle and distance difference values relating for each pair. **g**–**h**, Separately linearizing (**g**) and the angle and distance differences between all pairs and then concatenating them (**h**) enables visualization of changes in these positional arrangements across slices. **i**–**k**, Extension of 2D symmetry analysis of axon neighbour relations to the same set of 44 identified reticulospinal neurons that were analyzed in 3D. **i**, Top view of the 22 identified axon pairs for which 2D analysis was performed. **j**–**k**, Viewing linearized angle (**j**) and distance (**k**) differences for tested pairs shows a generally similar trend toward stable neighbour relations as the smaller subset analysis. Most neighbour relations return to a state of symmetry even after local perturbations, such as the entry of a new axon into the bundle. Black regions indicate where reconstruction data does not exist for a given pair. Scale bars: **a**,**d**,**i**, 50μm; **b**, 5μm.

**Extended Data Figure 10:**
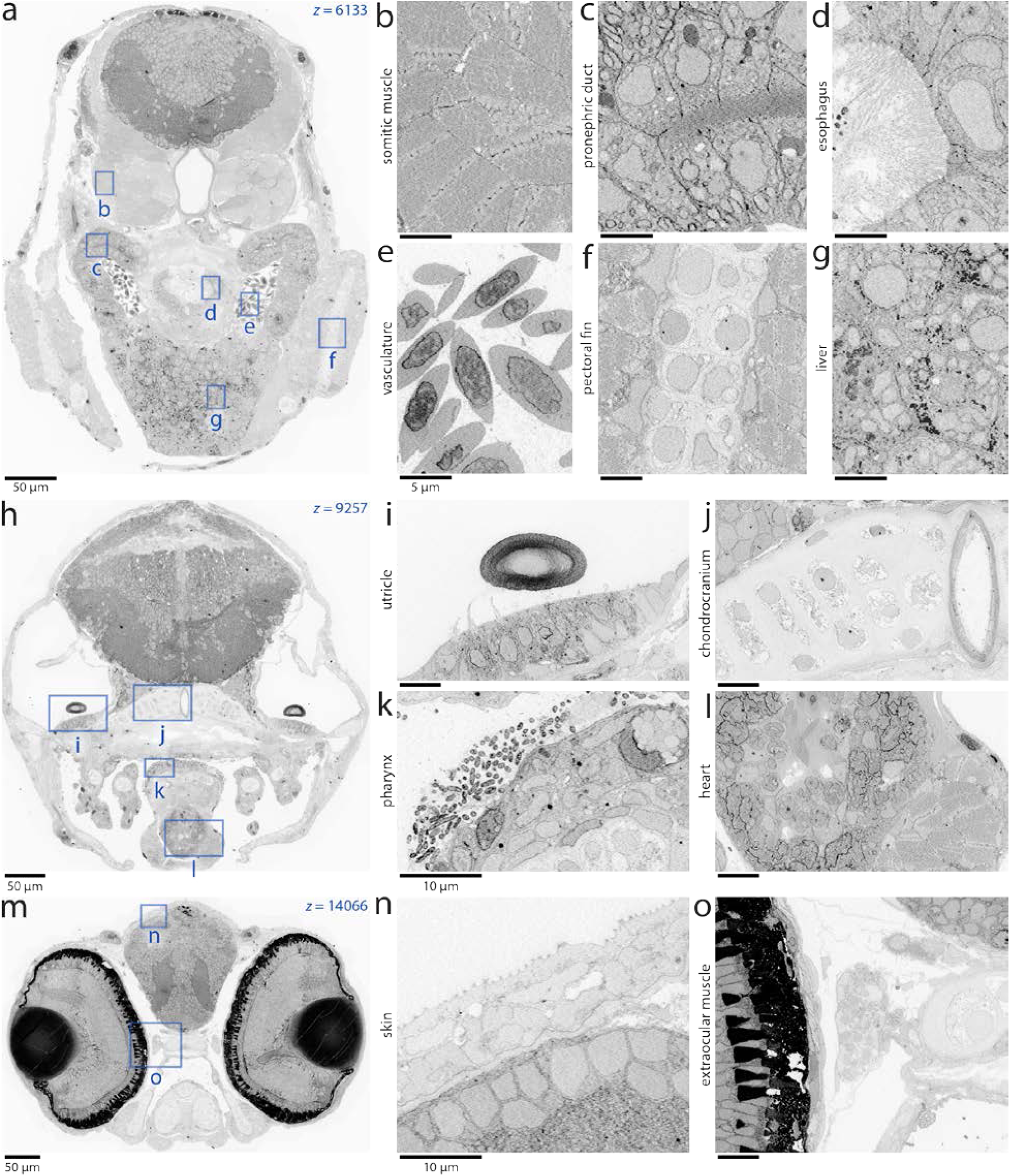
Examples of non-neuronal tissues contained within the dataset. In addition to capturing the whole brain, the nearly isotropic (56.4×56.4×60nm^3^vx^−1^) image volume containing the anterior quarter of a larval zebrafish serves as a high-resolution reference atlas for several other tissues and structures. Three selected sections (**a**,**h**,**m**) are accompanied by example images (**b–g**,**i–l**,**n–o**) to illustrate the variety of tissues and structures contained within the dataset. Scale bars: **a**,**h**,**m**, 50μm; **i**–**l**,**n**–**o**, 10μm; **b**–**g**, 5μm.

**Supplementary Video 1:** Survey of aligned section overview electron micrographs. Several sections excluded for faster viewing. Scale bar: 250μm.

**Supplementary Video 2:** Volume rendering of aligned section overview electron micrographs. Magenta and yellow planes correspond to reslice planes in Extended Data Figure 3c. Green plane corresponds to section outlined in Extended Data Figure 3a. Scale box: 250μm.

**Supplementary Video 3:** Survey of traverse larval zebrafish electron micrographs throughout the section series after image co-registration. Several sections excluded for faster viewing. Scale bar: 50μm.

**Supplementary Video 4:** Flythrough of serial micrographs centered on a reconstruction of the left Mauthner cell (magenta dot) beginning from its lateral dendrite, through the soma, through the axon cap, crossing to the contralateral side, and ending at its position in the spinal cord. Vertical line segments through these micrographs correspond to the contour reslices in Figure 1a-c. Several sections excluded for faster viewing. Scale bar: 5μm.

**Supplementary Video 5:** Matched horizontal reslice planes through the ssEM dataset and warped *in vivo* two-photon e*lavl3*:GCaMP5G fluorescence datasets. Scale bar: 100μm.

**Supplementary Video 6:** Serial electron micrographs centered on a reconstruction of a lateral line afferent neuron innervating the right dorsal (D2) neuromast (orange dot; Figure 3a-e). The video starts in high-resolution (4×4×60nm^3^vx^−1^) data of the neuromast where the afferent receives input from a hair cell (Figure 3b-c). The field-of-view then expands to follow the myelinated afferent in nearly isotropic resolution (56.4×56.4×60nm^3^vx^−1^) data throughout the periphery. It then magnifies to higher resolution (18.8×18.8×60nm^3^vx^−1^) data when approaching the afferent soma position in the peripheral lateral line ganglion before the afferent unmyelinates and enters the brain. Scale bar: 1μm (size changes depending on zoom in video).

**Supplementary Video 7:** Rotating volume rendering of the projectome consisting of myelinated axons manually reconstructed from the multi-resolution ssEM dataset. Colors are randomly assigned. Scale bar: 100μm.

**Supplementary Video 8:** Rotating rendering of lateral line system reconstructions revealed a striking bilateral symmetry. Posterior lateral line afferents are shown in yellow. Anterior and cranial lateral line afferents are shown in purple. Most neuromasts (blue) have a bilaterally symmetric counterpart except in one case (orange). Scale bar: 100μm.

**Supplementary Video 9:** Volume rendering of ssEM data and the Z-Brain spinal backfill label (red) combined with reticulospinal neuron reconstructions. This cross-modal registration of Z-Brain atlas data allowed for the anatomical characterization of domains within the ssEM dataset and aided with validation of neuron identification. Scale bar: 100μm.

**Supplementary Video 10:** Visualization of two-dimensional (2D) symmetry analysis of axon neighbour relations. Angle and distance differences (top right) were calculated from cross-sections (top left) of a subset of myelinated axon reconstructions (bottom), linearized, and aggregated (middle) for display. This video explains how the 2D symmetry analysis was performed across slices and illustrates how panels in Figure 4 and Extended Data Figure 9 were constructed.

